# Schizotypy-related magnetization of cortex in healthy adolescence is co-located with expression of schizophrenia-related genes

**DOI:** 10.1101/487108

**Authors:** Rafael Romero-Garcia, Jakob Seidlitz, Kirstie J Whitaker, Sarah E Morgan, Peter Fonagy, Raymond J Dolan, Peter B Jones, Ian M Goodyer, John Suckling, the NSPN Consortium, Petra E Vértes, Edward T Bullmore

**Author notes:** Correspondence to: Rafael Romero-Garcia PhD, University of Cambridge, Department of Psychiatry, Sir William Hardy Building, Downing Street, Cambridge CB2 3EB, UK. Full member list available in Supplemental Information (the NSPN Consortium). These authors contributed equally (Petra E Vértes, Edward T Bullmore).

## Abstract

**Background:** Genetic risk is thought to drive clinical variation on a spectrum of schizophrenia-like traits but the underlying changes in brain structure that mechanistically link genomic variation to schizotypal experience and behaviour are unclear.

**Methods:** We assessed schizotypy using a self-reported questionnaire, and measured magnetization transfer (MT), as a putative micro-structural MRI marker of intra-cortical myelination, in 68 brain regions, in 248 healthy young people (aged 14-25 years). We used normative adult brain gene expression data, and partial least squares (PLS) analysis, to find the weighted gene expression pattern that was most co-located with the cortical map of schizotypy-related magnetization (SRM).

**Results:** Magnetization was significantly correlated with schizotypy in bilateral posterior cingulate cortex and precuneus (and for disorganized schizotypy also in medial prefrontal cortex; all FDR-corrected *P* < 0.05), which are regions of the default mode network specialized for social and memory functions. The genes most positively weighted on the whole genome expression map co-located with SRM were enriched for genes that were significantly down-regulated in two prior case-control histological studies of brain gene expression in schizophrenia. Conversely, the most negatively weighted genes were enriched for genes that were transcriptionally up-regulated in schizophrenia. Positively weighted (down-regulated) genes were enriched for neuronal, specifically inter-neuronal, affiliations and coded a network of proteins comprising a few highly interactive “hubs” such as parvalbumin and calmodulin.

**Conclusions:** Microstructural MRI maps of intracortical magnetization can be linked to both the behavioural traits of schizotypy and to prior histological data on dysregulated gene expression in schizophrenia.

## Introduction

The genetic architecture of schizophrenia spectrum disorders assumes many independent allelic variations, each of small effect, contributing to the probability of diagnosis. Individuals with the greatest accumulation of genetic risk have the more severe psychotic disorder; individuals with a lower genetic risk may have less severe, non-psychotic schizotypal personality disorder (1), characterized by social eccentricity and unusual beliefs (2). The genetic risk for schizophrenia has been resolved more clearly by recent genome-wide association studies (3, 4) and by post-mortem human brain transcriptional studies (3, 5). However, it remains unclear how expression of these schizophrenia-related genes might be related to neuroimaging markers of schizophrenia spectrum disorders.

Macro-structural magnetic resonance imaging (MRI) studies – which measure anatomical parameters like cortical thickness – have collectively provided robust evidence for reduced volume or thickness in a network of inter-connected cortical areas in patients with schizophrenia (6). There have been fewer MRI studies of schizotypy and the pattern of macro-structural results has not been consistent, perhaps reflecting their relatively small sample sizes (**Table 1**).

**Table 1:** Previous studies relating schizotypal traits and macro-structural MRI metrics. Directionality refers to whether the study reports a positive (+) or negative (-) association between a schizotypy measure and a structural metric; VBM: Voxel-based Morphometry; R: Right; L: Left; CAPE: Community Assessment of Psychic Experiences; RISC: Rust Inventory of Schizotypal Cognitions; PSI-C: Psychiatric and Schizotypal Inventory For Children; FWE: family-wise error; SIS: Structured Interview for Schizotypy.

| Authors | Sample Size | Mean age (years) | Experimental Design | Brain coverage | Schizotypy measure | Type I error control | Structural index | Directionality | Brain regions |
| --- | --- | --- | --- | --- | --- | --- | --- | --- | --- |
| *Evans et al. 2016* [7] | 28 | 11 | VBM | Whole brain | PSI-C | Bonferroni | Volume | - | Caudate, amygdala, hippocampal gyrus, middle temporal |
| *Nenadic et al. 2015* [8] | 59 | 31 | VBM / Positive and negative factor of schizotypy | Whole brain | CAPE | FWE | Volume | - | R. precuneus |
| *Wang et al. 2015* [9] | 69 | 19 | VBM / Dividing subjects with high/low schizotypy | Whole brain | SPQ | AlphaSim permutation | Volume | - | Dorsolateral prefrontal cortex, insula, posterior temporal, cerebellum |
| *DeRosse et al. 2015* [10] | 138 | 36 | ANCOVA/ Dividing subjects with high/low schizotypy | ROIs | SPQ | None | Thickness / GM and WM Volume | - | Frontal, temporal |
| *Kuhn et al. 2012* [11] | 34 | 36 | Vertex-wise / Positive and negative factor of schizotypy | Whole cortex/ Thalamus | SPQ | FDR | Thickness / Thalamus volume | + | R. dorso lateral prefrontal cortex, R. dorsal premotor |
| *Ettinger et al. 2012* [12] | 55 | 27 | VBM | Whole brain | RISC | FWE cluster | Volume | - | Medial prefrontal, orbitofrontal, temporal |
| *Modinos et al. 2010* [13] | 38 | 20 | VBM/ Dividing subjects with high/low Schizotypy | Whole brain | CAPE | FDR | Volume | + | Medial posterior cingulate, precuneus |
| *Moorhead et al. 2009* [14] | 98 | 16 | VBM / Longitudinal study | Whole brain | SIS | Unspecified multiple comparison correction | Volume | - | L. medial temporal, L. amygdala, L. parahippocampal gyrus |
| *Stanfield et al. 2008* [15] | 143 | 16 | ANCOVA | Whole brain | SIS | None | Folding | + | R. prefrontal |

Micro-structural MRI provides information about the composition of tissue within a voxel (7). For example, magnetization transfer (MT) images (8), and “myelin maps” derived from the ratio of conventional T1- and T2-weighted images (T1w/T2w) (9), are sensitive to the proportion of fatty brain tissue represented by each voxel which, according to histological studies on animal models, is related to myelin content (10–13). MT maps have been used as markers of myelination, in white matter and in cortex (14) in healthy subjects (15) and in demyelinating disorders such as multiple sclerosis (16–18). Schizophrenia has been associated with reduced MT in frontal, temporal and insular cortex (19–26) and the cortical expression of schizophrenia-related genes was (negatively) correlated with T1w/T2w maps (27).

In this context, we measured schizotypy, using the Schizotypal Personality Questionnaire (SPQ), and magnetization transfer (MT), using a multiparameter MRI scanning procedure, in a sample of 248 healthy young people (14-25 years) (**Table S1**). We tested three key hypotheses in a logical sequence. (*i*) that intra-cortical MT was correlated with SPQ total score (and subscale scores), (*ii*) that the cortical pattern of schizotypy-related magnetization was co-located with a cortical map of weighted whole genome expression, and (*iii*) that the gene transcripts most strongly coupled to schizotypy-related magnetization were enriched for genes that were transcriptionally dysregulated in histological case-control studies of schizophrenia.

## Methods

### Sample

2135 healthy young people, aged 14-25 years, were recruited from schools, colleges, NHS primary care services and direct advertisement in north London and Cambridgeshire, UK. This primary cohort was stratified into 5 contiguous age-related strata, balanced for sex and ethnicity (28). A secondary cohort of N=297 was recruited by randomly sub-sampling the primary cohort so that ∼60 participants were assigned to each of the same age-related strata, balanced for sex and ethnicity, as in the primary cohort. Participants were excluded if they had a current or past history of clinical treatment for a psychiatric disorder, drug or alcohol dependence, neurological disorder including epilepsy, head injury causing loss of consciousness, or learning disability; see **Supplemental Information** (SI) for details.

Written informed consent was provided by all participants as well as written parental assent for participants less than 16 years old. The study was approved by the National Research Ethics Service and conducted in accordance with National Health Service research governance standards.

### Schizotypy assessment

The Schizotypal Personality Questionnaire (SPQ) (29) is a self-report scale, comprising 74 dichotomous items that are grouped on 9 subscales, measuring the complex trait of schizotypy. Participants completed the SPQ on two assessments, separated by 6-18 months, so that trait-like scores on total and subscale SPQ metrics could be estimated by the number of questionnaire items positively endorsed by each participant on average over time.

### MRI data acquisition

Structural MRI scans were acquired on one of three identical 3T MRI systems in London or Cambridge, UK (Magnetom TIM Trio, Siemens Healthcare, Erlangen, Germany; VB17 software version). The multi-parametric mapping (MPM) protocol (8) yielded 3 multi-echo fast low angle shot (FLASH) scans with variable excitation flip angles. By appropriate choice of repetition time (TR) and flip angle α, acquisitions were predominantly weighted by T1 (TR=18.7ms, α=20°), proton density (PD), or magnetization transfer (MT) (TR=23.7ms, α=6°). Other acquisition parameters were: 1 mm^3^ voxel resolution, 176 sagittal slices and field of view (FOV) = 256 × 240 mm. A pilot study demonstrated satisfactory levels of between-site reliability in MPM data acquisition (8). MT images (15) and T1 images (30, 31) from this sample have been previously reported.

### MRI reconstruction, cortical parcellation and estimation of schizotypy-related magnetization

We used a standard automated processing pipeline for skull stripping, tissue classification, surface extraction and cortical parcellation (http://surfer.nmr.mgn.harvard.edu) applied to longitudinal relaxation rate (R1) maps (R1=1/T1). Expert visual QC ensured accurate segmentation of pial and grey/white matter boundaries. Regional MT values were estimated at each of 68 cortical regions for each subject, resulting in a (248 × 68) regional MT data matrix. The Euler number for the R1 images was calculated as a proxy measure of image quality in the simultaneously acquired MT images (32).

A simple linear model of age-related change in MT was used to estimate two key parameters for each region: baseline MT at 14 years (MT_14_), and age-related rate of change in the period from 14 to 24 years old (ΔMT) (15). For the principal analyses, effects of age on MT were controlled by regression before estimating the correlation of the age-corrected MT residuals with total SPQ. The Kolmogorov-Smirnov normality test was used to determine the appropriate correlation estimator (Pearson’s or Spearman’s).

### Estimation of regional gene expression

We used the Allen Human Brain Atlas (AHBA), a whole genome expression atlas of the adult human brain created by the Allen Institute for Brain Sciences using six donors aged 24-57 years (http://human.brain-map.org) (33). Probe-to-gene and sample-to-region mapping strategies can have a major impact on regional gene expression estimation (34). Here we used the genome assembly hg19 (http://sourceforge.net/projects/reannotator/; (35)) to reannotate the probe sequences into genes (36). When genes were mapped by multiple cRNA hybridization probes, the probe showing highest average expression across samples was selected (37). MRI images of the AHBA donors were parcellated using the Desikan-Killiany atlas and each cortical tissue sample was assigned to an anatomical structure. Regional expression levels were compiled to form a (68 × 20,647) regional transcription matrix (38); see **SI**.

### Schizotypy-related magnetization and human brain gene expression

We used partial least squares (PLS) to analyse covariation between SRM and gene expression because it is technically well-suited to the high collinearity of the gene expression data (39, 40); and because PLS and the related multivariate method of canonical correlation analysis (CCA) have been extensively developed and used for neuroimaging and transcriptional data analysis (41–44). Specifically, we used PLS to analyse the relationship between the vector of 68 regional measures of SRM and the (68 × 20,647) matrix of 68 regional mRNA measurements for 20,647 genes (44). The first PLS component (PLS1) was defined as the weighted sum of whole genome expression that was most strongly correlated, or most closely co-located, with the anatomical map of SRM. Permutation testing based on spherical rotations or “spins” of the spatially correlated SRM map was used to test the null hypothesis that PLS1 explained no more covariance between SRM and whole genome expression than expected by chance (*P*_spin_) (31). Bootstrapping was used to estimate the variability of each gene’s positive or negative weight on PLS1 and we tested the null hypothesis of zero weight for each gene with false discovery rate (FDR) of 5% (42). The set of genes that were significantly (positively or negatively) weighted on PLS1 was called the SRM gene list or set.

### Enrichment analysis

We assigned a cellular affiliation score to each gene in the SRM gene list according to prior criteria for four cell types: neuron, astrocyte, microglia or oligodendroglia (45); and for a more fine-grained set of cell types (46) (**Table S2**). We used a data resampling procedure to test the null hypothesis that SRM genes were randomly assigned to different cell types.

We used two lists of genes that were differentially expressed, or transcriptionally dysregulated, in post-mortem brain tissue measurements of mRNA from case-control studies of schizophrenia: (*i*) the list of genes reported by Gandal *et al* (2018) as up-regulated (845) or down-regulated (1175) in prefrontal and parietal brain regions in schizophrenia (N=159); and (*ii*) the list of genes reported by Fromer *et al*. (2016) as up-regulated (304) or down-regulated (345) in the dorsolateral prefrontal cortex in schizophrenia (N=258). The two gene lists were partially overlapping **(Table S3)** and differential expression of all genes subsumed by the union of the two lists was strongly correlated between studies (*ρ* = 0.76, *P* < 10^−129^, **Figure S1**).

We used repeated random re-labelling of genes to test the null hypothesis that the SRM gene list included no more schizophrenia-related genes than expected by chance. We also applied the same resampling procedures to comparable prior data on differential gene expression from case-control studies of inflammatory bowel disease (IBD), bipolar disorder (BPD), major depressive disorder (MDD), and autism spectrum disorder (ASD) (Gandal et al., 2018).

## Results

### Sample characteristics

After quality control checks, complete, evaluable MRI and behavioural data were available for analysis on 248 participants: age 19.11 ± 2.93 years [mean ± standard deviation]; 123 (50%) female; 213 (86%) right-handed; IQ = 112.0 ±10.5; 214 (86%) white Caucasian, 10 Asian, 4 Black / African / Caribbean, 17 mixed, and 3 other ethnic groups (see **Table S1** for details).

### Schizotypy and magnetization transfer

Schizotypal personality scores in this healthy (non-psychotic) sample followed a positively skewed distribution (mean = 0.23, median = 0.20) that was normalized by square root transformation prior to statistical analysis. There was no significant effect of age *(R*^*2*^ < 10^−3^, *P*=0.69; **Figure S2**), gender *(R*^2^ < 10^−3^; *P*=0.77) or age-by-gender *(R*^2^ < 10^−3^; *P*=0.82) on SPQ total score or subscale scores.

Total SPQ score was modestly positively correlated with global magnetization transfer, estimated as the average MT over all 68 regions (*R*^*2*^ = 0.02; *P* = 0.015; **Figure S3**). Total SPQ was significantly correlated with age-corrected regional MT in 4 out of 68 regions individually tested (*R*^*2*^ > 0.04, *P* < 0.05, FDR corrected; **Figure 1A, Figure 1B** and **Table S4)**: the left isthmus cingulate, left posterior cingulate, left precuneus and right isthmus cingulate. These medial posterior cortical regions had high MT signals at age 14 (MT_14_) and relatively slow rates of increase in MT over the period 14-25 years (ΔMT) (**Figure 1C**).

**Figure 1.**
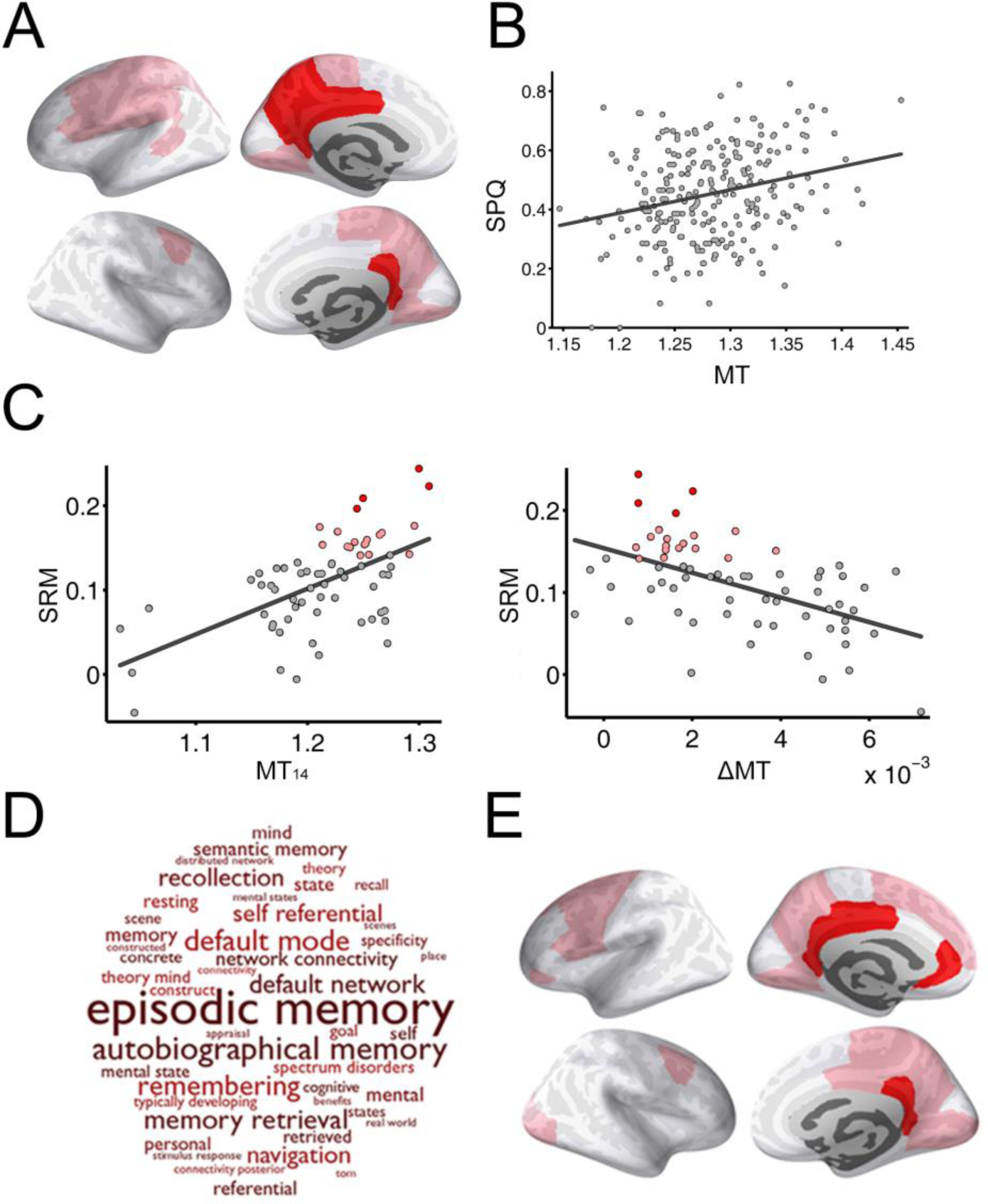
Schizotypy-related magnetization: association between intra-cortical magnetization transfer (MT) and schizotypal personality questionnaire (SPQ) score. (a) Cortical surface maps highlighting areas where total SPQ score was significantly positively correlated with regional MT after controlling for age by regression: pink regions had nominally significant schizotypy-related magnetization (SRM) (two-tailed *P* < 0.05); red regions had significant SRM controlled for multiple comparisons over 68 cortical regions tested (FDR < 0.05). (b) Scatterplot of SPQ total score for each participant versus mean MT in regions of significant schizotypy-related magnetization (*R*^*2*^ = 0.04, P = 0.002, df = 246); each dot represents one of 248 healthy people aged 14-25 years. (c) Scatterplots of SRM versus magnetization transfer at age 14 years (MT_14_) (left; *R*^*2*^ = 0.34, P_spin_ = 0.002, df = 67) and SRM versus change in magnetization aged 14-25 years (ΔMT) (right; *R*^*2*^ = 0.28, P_spin_ = 0.006, df = 67). Each point represents a cortical region and colored points represent regions with significant schizotypy-related magnetization (pink, *P* < 0.05; red, FDR < 0.05). (d) Word cloud representing ontological terms most frequently associated with functional activation of the medial posterior cortical areas of significant schizotypy-related magnetization. (e) Cortical surface maps highlighting areas where scores on the disorganized factor of schzioptypy was significantly positively correlated with regional MT after controlling for age by regression (pink, two-tailed *P* < 0.05; red, FDR < 0.05).

We used prior fMRI data to identify experimental task conditions that were most robustly associated with functional activation of these areas of significant schizotypy-related magnetization (http://neurosynth.org; (47)): memory, social cognition or theory of mind, and executive functions (**Figure 1D**). The posterior cingulate and medial parietal cortical areas of significant SRM were also enriched for default mode network-related terms in ontological analysis of prior fMRI data (47).

### Sensitivity analyses of schizotypy-related magnetization

We used a linear model to control the association between SPQ and MT for the potentially confounding effects of age, gender, site, socio-economic status and total brain volume (**Figure S4)**, and robust estimators to mitigate the influence of the small number of high SPQ scores on the estimation of SRM (**Figure S5)**. In both cases, the key results were conserved: namely, significant SRM in DMN areas and significant correlations between PLS1 weights and differential gene expression in schizophrenia. We also noted a negative correlation between Euler number and global MT, indicating reduced MT in a minority of poor quality images (*ρ* = −0.14, *P* = 0.03). When we excluded the 10% of participants with poorest image quality, the correlation between Euler number and MT was no longer significant (*ρ* = 0.04, *P* = 0.11) but the key results were conserved (**Figure S6)**.

Schizotypy is a complex trait comprising multiple dimensions of cognition, emotion and behaviour. In addition to the principal analysis of total SPQ, we also considered two possible decompositions of the schizotypal trait. Nine subscales of the SPQ defined by (29) were positively correlated with regional MT but these associations were less robust than for total SPQ (**Figure S7)**. All three factors of schizotypy defined by (48), i.e., positive, negative and disorganised dimensions, were positively correlated with MT. The correlation between disorganised schizotypy and MT was strongest and statistically significant controlling for multiple comparisons (**Figure 1E and Figure S8**).

We also measured cortical thickness (CT) for each of the same 68 regions, using R1 images collected as part of the same MRI sequence used to measure MT. SPQ scores were negatively correlated with cortical thickness in some regions but the associations between schizotypy and CT were not significant when corrected for multiple tests (**Figure S9)**.

### Schizotypy-related magnetization and gene expression

The first partial least squares component (PLS1) defined a weighted sum of whole genome expression that accounted for ∼40% of the cortical patterning of schizotypy-related magnetization, significantly more than expected by chance (*P*_*spin*_=0.027; **Figure 2A**).

**Figure 2.**
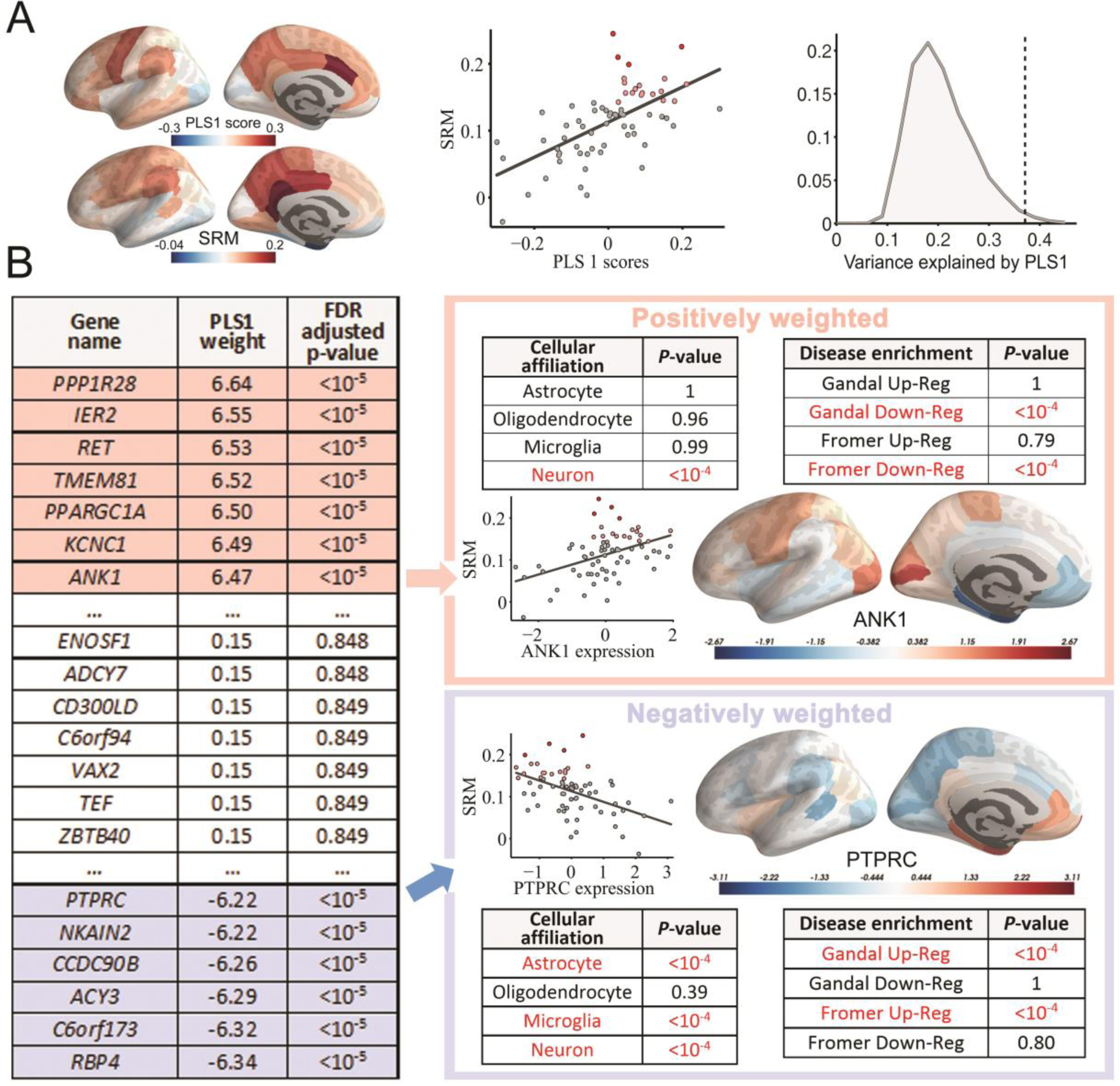
Gene expression and schizotypy-related magnetization. (a) (left) The first partial least squares component (PLS1) defined a linear combination of genes that had a similar cortical pattern of expression to the cortical map of schizotypy-related magnetization, representing the correlation between SPQ and MT at each of 68 cortical regions. (center) Scatterplot of PLS1 scores versus schizotypy-related magnetization; each point is a cortical region. (right) The combination of genes defined by PLS1 explains more of the variance in schizotypy-related magnetization (dotted line) than expected by chance (histogram of permutation distribution) (b) Illustrative example of the weights assigned to representative genes on PLS1. Genes with the highest positive weights are colored in pink, non-significantly weighted genes are shown in white, and the genes with the lowest negative weights are colored in blue. Tables summarise *P*-values by permutation testing for enrichment analysis by four lists of genes affiliated to specific cell types and four lists of genes associated with schizophrenia: Gandal and Fromer up-reg are lists of genes transcriptionally up-regulated post-mortem in schizophrenia; Gandal and Fromer down-reg are lists of genes transcriptionally down-regulated in schizophrenia. Scatterplots and cortical maps illustrate that positively-weighted genes, like *ANK1*, are over-expressed in cortical regions with high levels of schizotypy-related myelination; whereas negatively-weighted genes, like *PTPRC*, are under-expressed in regions with high levels of SRM.

Multiple univariate *Z*-tests were used to test the set of null hypotheses that the weight of each gene on PLS1 was zero. We found that this null hypothesis was refuted for 1,932 positively-weighted genes and for 2,153 negatively-weighted genes (*P* < 0.05, FDR corrected for whole genome testing at 20,647 genes; **Figure 2B**). Positively-weighted genes were normally over-expressed, and negatively-weighted genes were normally under-expressed, in cortical areas with high schizotypy-related magnetization. These 4,085 genes constituted the schizotypy-related magnetization (SRM) gene list.

### Functional and schizophrenia-related enrichment of schizotypy-related magnetization genes

The SRM gene list was tested for enrichment by genes characteristic of specific cell types using two sets of prior criteria (45, 46). Positively-weighted SRM genes were enriched for neuronal affiliation (45) (permutation test, *P* < 10^−4^) and, more specifically, for genes differentially expressed in fast-spiking parvalbumin-positive inhibitory interneurons (46) (**Table S2**; permutation test, FDR-corrected *P* < 0.01). Negatively-weighted SRM genes were enriched for astrocytes, microglia and neuronal affiliation (45, 46) (permutation tests, all P < 10^−4^) (**Figure 2B)**.

The positive or negative weighting of each SRM gene was strongly related to its differential expression in two post-mortem studies of schizophrenia (49). Positively-weighted SRM genes were enriched for genes that were significantly down-regulated in both studies (**Table S5**); but not for significantly up-regulated genes in either study. Additionally, positively-weighted SRM genes were also enriched for genes previously associated with white-matter connectivity differences in schizophrenia described by (58) (**Figure S10**). In contrast, negatively-weighted SRM genes were enriched for genes that were significantly up-regulated in both studies (permutation tests, all *P* < 10^−4^) (**Figure 2B**) (**Table S6**); but not for significantly down-regulated genes.

Convergently, there was a significant negative correlation (Spearman’s *ρ* = −0.16, *P* < 10^−6^) between the PLS1 weights of all genes in the genome and the differential expression values reported for all genes by Gandal et al (2018) and Fromer et al. (2016) (**Figure 3A**). PLS1 gene weights were not correlated with differential expression in inflammatory bowel disease or major depressive disorder; however, they were negatively correlated with differential expression in bipolar disorder and autism spectrum disorder (**Figure 3B**).

**Figure 3.**
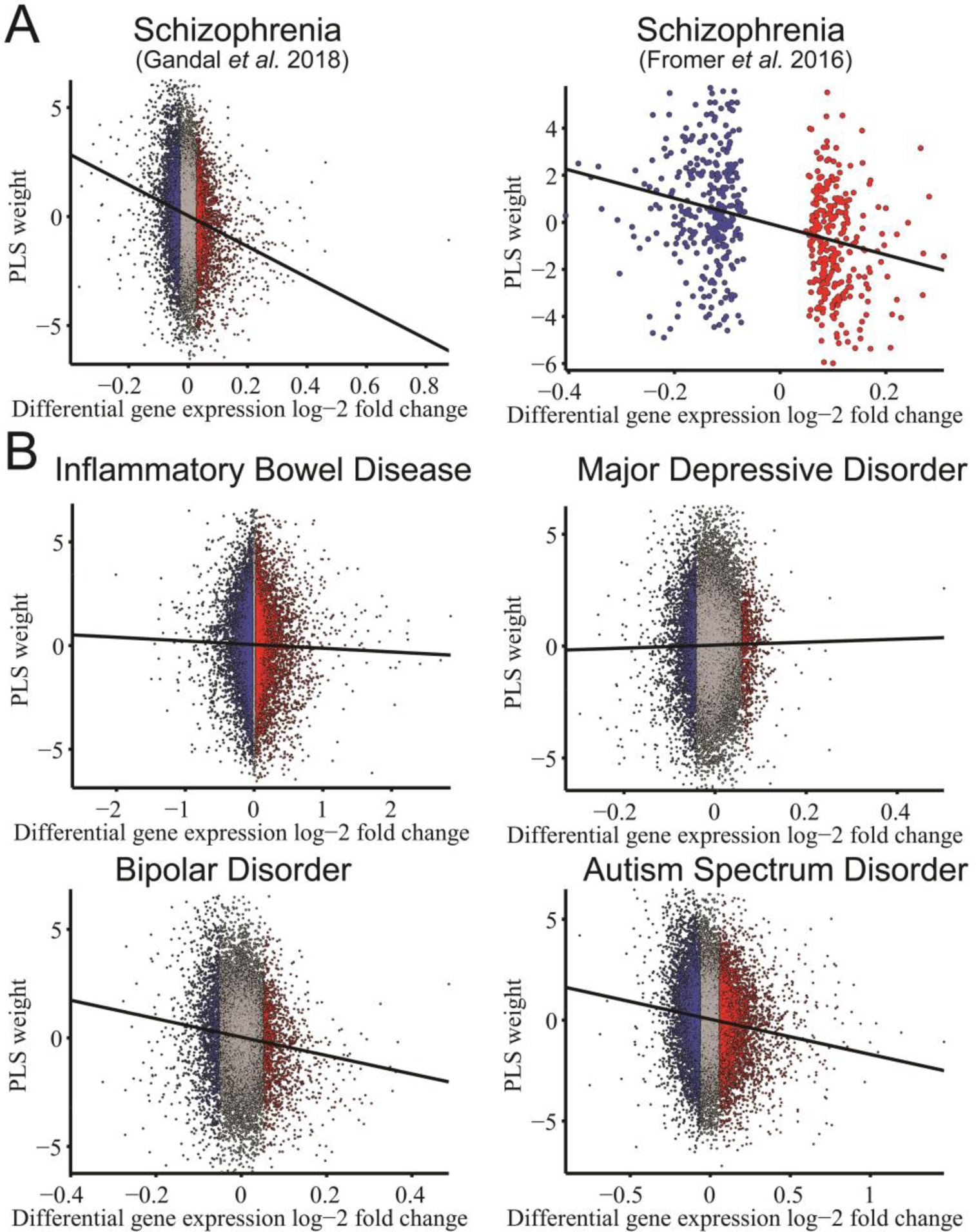
Weights of gene expression from PLS analysis of schizotypy-related magnetization were related to histological measures of differential gene expression from case-control studies of schizophrenia and other disorders. (a) The weight of each gene on the first PLS component was significantly negatively correlated with differential gene expression post-mortem in schizophrenia according to prior data reported by Gandal et al (2018) (Spearman’s rank correlation, *ρ* = −0.16, Bonferroni-corrected P_adj_ < 10^−65^, df = 11111) and by Fromer et al (2016) (Spearman’s rank correlation, *ρ* = −0.30, P_adj_ < 10^−12^, df = 586; for this dataset only significantly different expression values have been reported (46)). (b) Correlations between PLS weights and differential expression were also evaluated for other conditions (5): inflammatory bowel disease (*ρ* = −0.02, P_adj_ = 0.10, df = 586), major depressive disorder (*ρ* = 0.007, P_adj_ = 0.37, df = 15281), bipolar disorder (*ρ* = −0.09, P_adj_ < 10^−19^, df = 16064) and autism spectrum disorder (*ρ* = 0.11, P_adj_ < 10^−35^, df = 11131). Red and blue points represent genes that are significantly up- and down-regulated in post-mortem data.

We analysed the network of known protein-protein interactions (PPI) (STRING (50); http://string-db.org) between proteins coded by the 213 genes that were significantly down-regulated in schizophrenia (Gandal *et al* 2018) and significantly positively-weighted in the PLS analysis of schizotypy-related magnetization (**Table S5**). There were significantly more interactions (edges) between proteins coded by these genes than expected by chance (**Figure 4 and S11**; permutation test, *P*<10^−5^). Topologically, the network comprised several clusters of densely interconnected and functionally specialised proteins. The biggest cluster was enriched for synaptic terms and centred around highly connected “hub” proteins (**Figure 4**).

**Figure 4.**
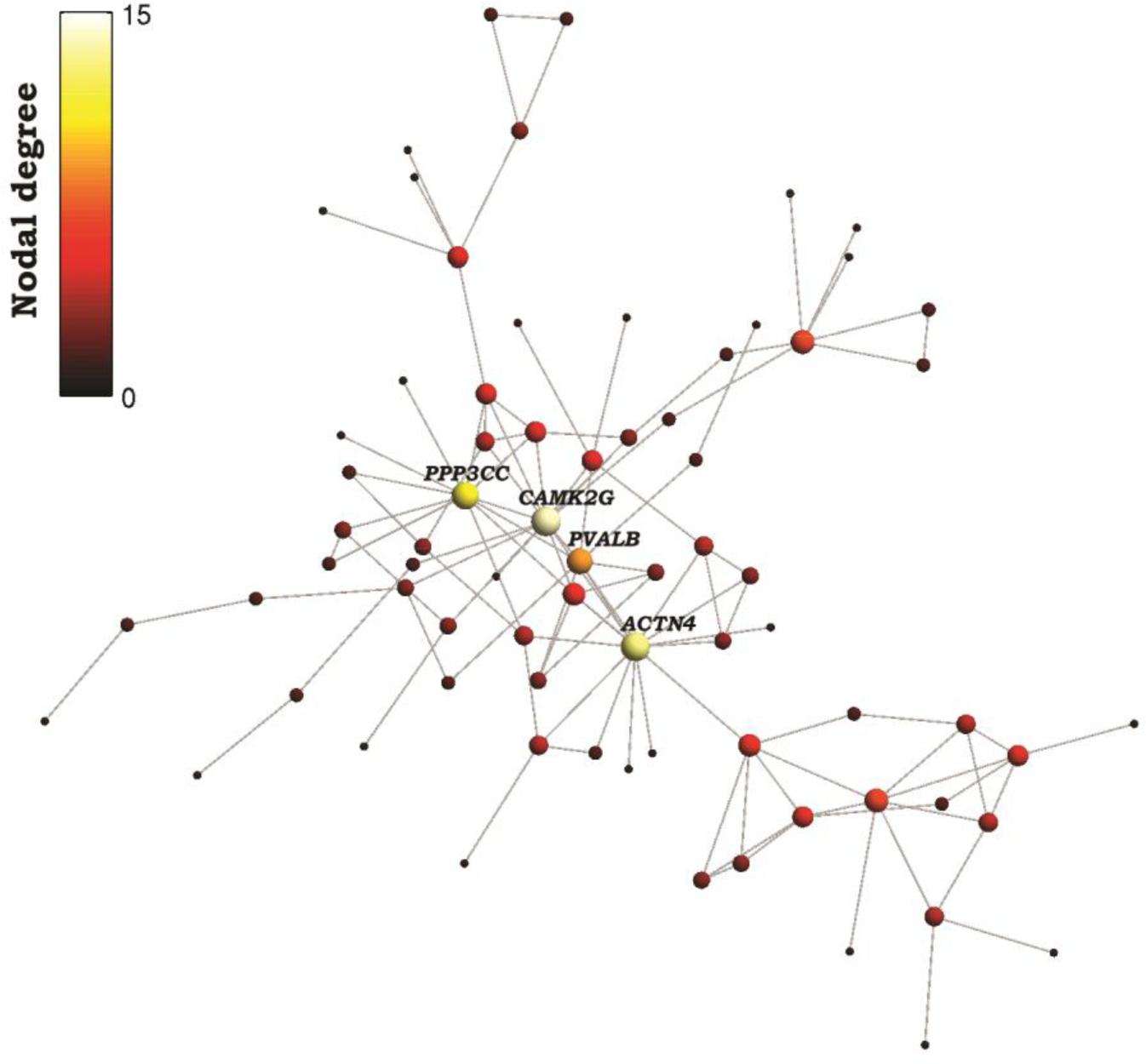
Protein-protein interaction network for a set of 213 proteins coded by genes associated with both schizotypy-related magnetization and post-mortem brain transcriptional dysregulation in schizophrenia. Nodes represent genes that were both (i) down-regulated in brain tissue from 159 patients with schizophrenia; and (ii) positively-weighted on the PLS component most strongly associated with schizotypy-related myelination in 248 healthy adolescents. Edges represent known protein-protein interactions. Color and size of each node represents its degree centrality or “hubness”, simply the number of interactions that protein has with the other proteins in the network. The top four most highly connected hubs are highlighted: PPP3CC is a calmodulin dependent phosphatase, calcineurin; CAMK2G is a calcium/calmodulin dependent kinase; PVALB is a calcium binding protein, parvalbumin; ACTN4 is a microfilamentous protein, actinin alpha 4. This network is specialised for calcium dependent processes that have been previously associated with interneurons and with pathogenesis of schizophrenia. For the complete list of gene names on the PPI network see **Figure S11** and go to https://version-10-5.string-db.org/cgi/network.pl?taskId=RMpA04wbWG8k for a full interactive version of the PPI network.

## Discussion

Schizotypy was associated with intra-cortical magnetization of posterior cortical regions of the default mode network and co-located with a normative cortical pattern of weighted whole genome expression. The gene transcripts most strongly weighted in association with this MRI map of schizotypy-related magnetization (SRM) were significantly enriched for genes specifically expressed by neuronal and glial cells, especially parvalbumin-positive interneurons, and for genes that were transcriptionally dysregulated in two prior post-mortem studies of schizophrenia.

### Magnetization, intra-cortical myelination and schizotypy

Magnetization transfer is a micro-structural MRI measurement that is sensitive to the ratio of fatty and watery tissue represented by each voxel and, in the brain, most of the fat is myelin. Intra-cortical myelination, especially of the deeper layers of cortex, has been recognised since seminal cytoarchitectonic and myeloarchitectonic studies in the early 20^th^ century (14). MT images represent a stark contrast between the cortex and the central white matter as well as more nuanced variations across different cortical areas and layers (51). Histological measurements of myelin were positively correlated with MRI measurements of MT in post-mortem brains (52). Intra-cortical measurements of MT in humans have been validated as micro-structural MRI markers of myelination in healthy volunteers (53) and in patients with multiple sclerosis (54).

One plausible interpretation of schizotypy-related magnetization, therefore, is as a proxy marker for a biological state of schizotypy-related myelination. On this assumption, the results are open to further interpretation at a cellular level. For example, greater “myelination” could imply a greater density of myelinated neurons per voxel (a neuronal process), or a greater density of myelin per neuron (an oligodendroglial process), or some combination of these and other cellular parameters. The data available to us did not allow direct resolution of the relationships between schizotypy-related magnetization and myelination. Instead, we used open data on human brain gene expression (N=6, mean age = 42.5 years) to explore these questions more indirectly.

The adult brain gene expression profile that was most closely co-located with the adolescent brain map of SRM (N=248, mean age = 19) was enriched for neuronal, but not oligodendroglial, affiliations. This pattern of results arguably favours the interpretation that higher MT indicates a greater density of myelinated neurons per voxel, rather than a greater density of myelin per neuron, in people with higher schizotypy scores. However, the 20+ year age gap between the MRI measurements and the mRNA measurements precludes definitive resolution of these and other possible cellular interpretations of schizotypy-related magnetization. Although the SRM gene set is not known to demonstrate major developmental changes in expression after childhood (**Figure S12)**, in future it will be important to co-locate MT phenotypes in children and young people (and animal models) with more precisely age-matched data on brain gene expression and histology.

The macroscopic medial posterior cortical areas where MT was most strongly correlated with schizotypy in general – total SPQ - are key components of the default mode network (as defined by functional MRI studies (55)) and specialised for memory, social cognitive and theory-of-mind functions that are known to be abnormal in patients with schizophrenia (56). Interestingly, the schizotypal factor of disorganization was also correlated with MT in medial prefrontal cortical areas that also form part of the DMN. All these regions had high levels of magnetization at the age of 14 years and no significant subsequent change in magnetization over the period 14-25 years. This contrasts with areas of lateral association cortex, which have a low level of MT at 14 years but significant increase in MT over the course of adolescence (15). We can infer that these medial posterior cortical areas matured as part of a pre-adolescent wave of cortical development (57), which would be compatible with the stable, trait-like properties of schizotypal personality in these data and in other studies of adolescents and adults.

### Cortical gene expression, schizotypy-related myelination and schizophrenia

We wanted to identify which genes in the whole genome had a cortical expression pattern that was most similar to the cortical map of schizotypy-related magnetization. A large number (>20,000) of non-independent statistical tests would be entailed in testing the association between each transcript’s spatially correlated cortical expression map and the cortical map of SRM. Therefore we favoured a multivariate approach and used partial least squares to identify a cortical pattern of weighted whole genome expression that was significantly co-located with the SRM map, and to identify which particular gene transcripts were most positively or negatively-weighted.

We found that the positively-weighted genes (1,932) were over-expressed in cortical areas with high levels of SRM; whereas the negatively-weighted genes (2,153) were over-expressed in cortical areas with low levels of SRM. Both positive and negative genes were enriched for neuronal, but not for oligodendroglial, affiliations. Positive genes were specifically enriched for parvalbumin positive inhibitory interneurons; negative genes were enriched for astrocytes and microglia.

We predicted hypothetically that the SRM gene set would be enriched for genes that are known to be transcriptionally dysregulated in schizophrenia. This prediction was supported by convergent results from enrichment analysis using two prior, independently discovered, and partially overlapping lists of genes differentially expressed in post-mortem case-control studies of schizophrenia. In both cases, there were significantly more histologically down-regulated genes in the list of positively-weighted SRM genes, and significantly more up-regulated genes in the list of negatively-weighted SRM genes, than expected by chance. In other words, genes with reduced brain transcription post-mortem in schizophrenia were normally more highly expressed in cortical areas with higher levels of schizotypy-related magnetization. A subset of the positively-weighted SRM genes has been previously associated with white-matter dysconnectivity in schizophrenia (58), suggesting that schizotypy related magnetization of cortex and schizophrenia-related disruption of central white matter tracts may be different imaging phenotypes related to expression of genes in common.

The sub-set of 213 SRM positive genes that were also significantly down-regulated in one or both of the prior histological studies coded for a protein-protein interaction (PPI) network comprising a small number of highly-connected hub proteins (ACTN4, CAMK2G, PPP3CC and PVALB), each hub having up to 14 known biochemical interactions with other proteins in the network. Calcium/calmodulin dependent protein kinase II gamma (CAMK2G) is one of a family of serine/threonine kinases that mediate many of the second messenger effects of Ca^2+^ that are crucial for plasticity at glutamatergic synapses. Parvalbumin (PVALB) is a calcium-binding albumin protein that is expressed particularly by the fast-spiking class of GABAergic interneurons that has been strongly implicated in the pathogenesis of schizophrenia (59) (**Figure 4**).

We found that the SRM gene list was also enriched by genes differentially expressed in autism spectrum and bipolar disorders (ASD and BPD). These results are consistent with the post mortem evidence (5) that the differential gene expression profile of schizophrenia (compared to healthy controls) is strongly correlated with the case-control differences of transcription in BPD and ASD (*ρ* > 0.45, *P* < 0.001). They are also consistent with clinical evidence that ASD and BPD are both associated with increased schizotypal traits (60, 61).

### Methodological issues

The brain tissue samples used for RNA sequencing in the AHBA were not homogeneously distributed across the cortex, so estimates of regional expression are based on different numbers of experimental measurements in each of the 68 regions. The case-control differences in frontal or parietal lobar gene transcription reported by Gandal et al (2018) and Fromer et al (2016), although based on a relatively large number of patients, are not as precisely localised or representative of the whole brain as the AHBA and MRI data. This study has a considerably larger sample size than any previously reported MRI study of schizotypy, and it is the first to evaluate a micro-structural MRI marker, which was more strongly related to schizotypy than the more conventional macro-structural MRI marker of cortical thickness. Nonetheless, it is theoretically surprising that there was limited evidence for significant schizotypy-related magnetization of frontal and lateral temporal cortex (although magnetization of medial prefrontal cortex was significantly associated with the disorganized component of schizotypy; **Figure 1E**), possibly reflecting limited statistical power. The SPQ is a self-report questionnaire measure of schizotypy; more refined and objective assessments of schizotypal traits would likely add value to future studies.

### Conclusions

Overall, these correlational results do not unambiguously resolve questions of causality but they are consistent with the interpretation that schizotypy-related magnetization, putatively an imaging marker of intra-cortical density of myelinated neurons, represents cellular processes determined in part by transcription of genes related to schizophrenia and other neuropsychiatric disorders.

## ACKNOWLEDGEMENTS

We thank the Allen Brain Institute for access to human brain gene expression data, as well as all the members of the NSPN consortium for data collection, storage and preprocessing: Edward Bullmore (Chief Investigator), Raymond Dolan, Ian Goodyer, Peter Fonagy, Peter Jones, Matilde Vaghi, Michael Moutoussis, Tobias Hauser, Sharon Neufeld, Michelle St Clair, Petra Vértes, Kirstie Whitaker, Rafael Romero-Garcia, Becky Inkster, Gita Prabhu, Cinly Ooi, Umar Toseeb, Barry Widmer, Junaid Bhatti, Laura Villis, Ayesha Alrumaithi, Sarah Birt, Aislinn Bowler, Kalia Cleridou, Hina Dadabhoy, Emma Davies, Ashlyn Firkins, Sian Granville, Elizabeth Harding, Alexandra Hopkins, Daniel Isaacs, Janchai King, Danae Kokorikou, Christina Maurice, Cleo McIntosh, Jessica Memarzia, Harriet Mills, Ciara O’Donnell, Sara Pantaleone, Jenny Scott, Pasco Fearon, Anne-Laura van Harmelen and Rogier Kievit. A previous version of this paper was posted on bioRxiv: https://www.biorxiv.org/content/10.1101/487108v1

## DATA and CODE SHARING

### Data

Regional magnetisation transfer (MT) for 68 cortical regions, schizotypy scores, age, gender, socio-economic status, scanning site, total brain volume and Euler values, for N=248, are available at: https://github.com/RafaelRomeroGarcia/Schizotypy_MT_geneExp.

### Code

Cortical parcellation of gene expression maps to estimate regional mean gene expression (62): (https://github.com/RafaelRomeroGarcia/geneExpression_Repository)

Generate null-models that preserve the spatial contiguity of cortical regions for permutation testing (31): https://github.com/frantisekvasa/rotate_parcellation

PLS analysis and bootstrapping to estimate PLS weights (15): https://github.com/KirstieJane/NSPN_WhitakerVertes_PNAS2016/tree/master/SCRIPTS

Generate Figure S1 from raw Gandal (5) and Fromer (49) datasets: https://github.com/RafaelRomeroGarcia/Schizotypy_MT_geneExp

Mapping regional values to the cortical surface for visualization (BrainsForPublication v0.2.1): https://doi.org/10.5281/zenodo.1069156

