## Supplementary Information for "Schizotypy-related magnetization of cortex in healthy adolescence is co-located with expression of schizophrenia-related genes"

***** These authors contributed equally

### Supplementary Methods

#### Sample

All primary cohort participants satisfied the following eligibility criteria: aged between 14 and 24 years inclusive; able to understand written and spoken English; willing and able to give informed consent for recruitment into the study cohort and consent to be re-contacted directly for possible participation in future studies within the consortium. They were excluded if they were currently or had recently (within the last 12 months) participated in a clinical trial of an investigational medical product. Participants completed questionnaire measures of socio-demographic status, family and educational or occupational environments, and sub-clinical psychopathology. All participants in the secondary MRI cohort satisfied the following additional eligibility criteria: willing and able to give informed consent for participation in the study and to attend full-day assessments at either the University of Cambridge or University College London study sites; have normal or corrected-to-normal vision. Participants were excluded if they had a current or past history of clinical treatment for a psychiatric disorder or for drug or alcohol dependence; had a current or past history of neurological disorders or trauma including epilepsy, or head injury causing loss of consciousness; had a learning disability requiring specialist educational support and/or medical treatment; or had a safety contraindication on the MRI scanning checklist. Secondary cohort participants attended one of the study sites (University of Cambridge or University College London) for a day of clinical, cognitive and MRI assessments. Socio-economic status was derived from the neighbourhood poverty index (NPI), provided by the UK Office of National Statistics (1).

#### MRI reconstruction, cortical parcellation and structural feature extraction

Freesurfer (recon-all) reconstructions were created for each participant using the longitudinal relaxation rate (R1) maps (R1=1/T1). Cortical thickness (CT) measurements were estimated by reconstructing the pial surface and the boundary between grey matter and white matter (2, 3) and measuring the distance between these surfaces. For each subject, MT-weighted images were projected on the cortical surface of each Freesurfer reconstruction (mri_vol2surf). CT and MT values were averaged across all vertices included in each of the 68 cortical parcel defined in the Desikan-Killiany atlas (4). This process was repeated for each subject resulting in a (248 × 68) matrix of cortical MT and a (248 × 68) matrix of CT at each of 68 regions for each of 248 participants.

*Meta-analysis of structural maps using Neurosynth*

To identify the cognitive functions served by the medial posterior cortical regions showing a significant correlation between MT and SPQ, a meta-analysis was performed using the Neurosynth software package (<http://neurosynth.org>; (5)). Neurosynth uses a combination of text mining, meta-analysis and machine-learning techniques to aggregate coactivation results from 11,406 functional MRI (fMRI) studies and links particular brain regions with particular cognitive functions based on keywords from the functional studies (5). This software was used here to find keywords associated with prior published findings correlated with the schizotypy-related myelination map. Neurosynth calculated the likelihood of each term given the activation described by the schizotypy-related myelination map -*i.e.* P(term | Activation map)-. The top 100 keywords (excluding anatomical terms for brain regions) ranked by the correlation strength (semantic association) between the MRI map and the meta-analytic map were visualized as a word cloud, with the size of the font scaled by correlation strength. This allowed us to identify the most likely reverse inferences that would be drawn if the schizotypy-related myelination map were a functional imaging map.

#### Estimation of regional gene expression

We used the Allen Human Brain Atlas (AHBA) – a whole-genome, whole-brain transcriptomic dataset of the adult human brain created by the Allen Institute for Brain Sciences (<http://human.brain-map.org>) (6). The AHBA includes six donors, three Caucasian, two African-American and one Hispanic. Their ages are 57, 55, 49, 39, 31 and 24 years. Further details on microarray analysis are available from the Allen Institute ([www.brain-map.org](http://www.brain-map.org)). According to AHBA annotation, 48171 out of 58692 probes can be mapped onto genes. Here, we used Richiardi et al.’s (2015) criterion to reannotate the probe sequences using the genome assembly hg19 (UCSC Genome Browser; <http://sourceforge.net/projects/reannotator/>; (7)). After the reannotation, a ﬁnal set of 49770 probes was associated with genes. Previous studies have shown the high between-study consistency of this probe-to-gene mapping method (8). Generally, multiple cRNA hybridization probes were used for each gene and, in these cases, we selected the probe with the highest mean gene expression. This is in line with recommendations from a recent study evaluating the impact of various preprocessing decisions on the correspondence between microarray and RNA-seq expression (9). Probes that were not matched to gene symbols in the AHBA data were excluded, resulting in 20,467 gene expression values that were evaluated in 3,702 brain samples. Due to the similarity of gene expression between hemispheres, the AHBA only sampled two of the four donors in the right hemisphere. In order to increase the number of AHBA samples per cortical region, all samples were pooled between bilaterally homologous cortical areas.

The anatomical structure associated with each tissue sample was determined using the MRI data provided by the AHBA for each donor. T1-images of the six donors were processed following the Freesurfer pipeline. Briefly, cortical surface reconstruction of each donor performed by Freesurfer recon-all involves (*i*) non-uniformity intensity correction, (*ii*) resampling to isotropic images of 1mm, *(iii)* registration to Talairach space, (*iv*) an initial WM segmentation that is used to generate a tessellated representation of the WM/GM boundary, (*v*) deformation of the tessellated WM/GM boundary outwards to the volume that maximizes the CSF/GM contrast to generate the pial surface and (*v*i) automatic correction of geometrical topological abnormalities. The Desikan-Killiany atlas employed in the neuroimaging dataset was reconstructed in each AHBA donor brain. Surface-based parcellation of each donor's brain was transformed into a volumetric parcellation that covered the whole cortical mantle. 73% of the total AHBA cortical samples were located within the resulting volumetric parcels. To improve on this, unassigned samples located less than 2 mm to a region boundary were additionally included to avoid exclusions due to registration misalignment, resulting in a final AHBA sample coverage of 91%. Gene expression values for each cortical parcel were estimated as the median gene expression of all the samples assigned to the parcel from all six donors. Expression values had very different scales across genes, so each gene expression value was normalized by taking a *Z*-score across regions. The normalised gene expression values were compiled as a (68 × 20,467) matrix that comprised transcriptional measurements for all 20,467 genes in the genome at each of 68 cortical regions.

#### Partial Least Squares analysis

Despite the high dimensionality on the transcriptomic data it is important to note that many of the genes have correlated expression patterns and this correlation structure severely reduces the effective degrees of freedom. To understand this, one may consider the extreme example where the dataset would consist of 20,000 identical (or perfectly correlated) genes – this would naturally be unlikely to reconstruct the 68-dimensional SRM map very well. In reality, the transcriptomic data are not so tightly (but still very highly) correlated. We note that this multicollinearity can be problematic for some statistical methods but the Partial Least Squares (PLS) method we use here was selected specifically for its ability to handle large datasets with high degrees on collinearity.

### Supplementary Results

#### Protein-protein interaction network enrichment

We used the software STRING (version 10.5) to plot the protein-protein interaction network amongst the 213 genes of interest and to overlay gene set enrichments onto the resulting network (<https://version-10-5.string-db.org/cgi/network.pl?taskId=RMpA04wbWG8k>). Figure S11 shows enriched GO Cellular Component terms and highlights in colour individual genes/proteins from the PPI network which are implicated in these enriched GO terms. The significantly enriched terms (after FDR correction) include some general terms such as “cytoplasm” or “cell part” and neuron-specific terms such as “synapse”, “neuron part” and “myelin sheath”. A similar analysis for significantly enriched GO Biological Process terms includes again general terms such as “transport” and terms more specific to neuronal function such as “synaptic transmission”.

#### Replication using only left hemisphere data

The gene expression dataset provided by AHBA includes two subjects with data across both hemispheres and four subjects with data only measured in the left hemisphere. This sampling strategy was chosen due to the similarity of gene expression between hemispheres. By mirroring the samples across hemispheres, we artificially duplicated the gene expression data. When we restricted the schizophrenia-enrichment analysis only to the left hemisphere, we found that positively weighted genes on PLS were significantly enriched for down-regulated genes (Gandal, P<10^-4^; Fromer, P=0.004) and negatively weighted genes on PLS were significantly enriched for up-regulated genes according to Gandal (P<10^-4^) but only a trend was observed for Fromer (P=0.059).

### Supplementary Figures

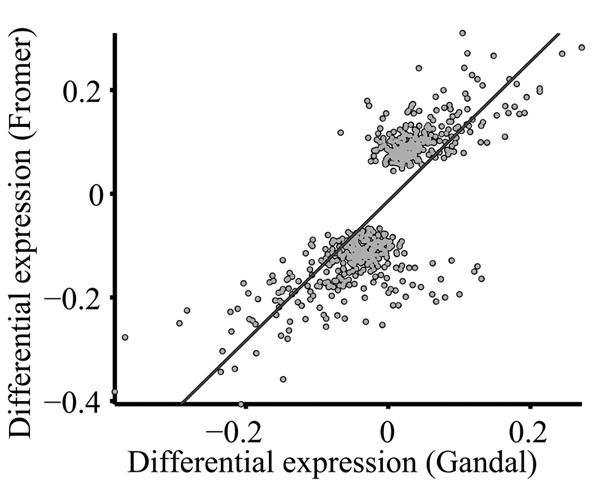

**Figure S1. Correlation between differential *post-mortem* gene expression in schizophrenia defined by Gandal and Fromer** (*ρ* = 0.76, *P* < 10^-129^, df=641). Script used to generate this figure is available at: <https://github.com/RafaelRomeroGarcia/Schizotypy_MT_geneExp/tree/master/Code_for_Figure_S1>

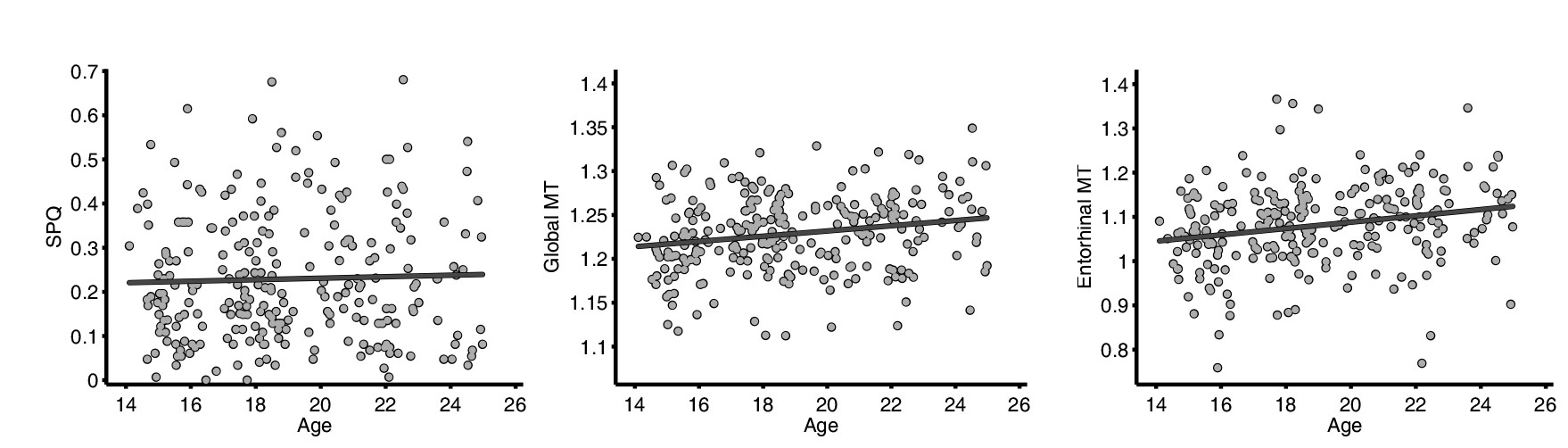

**Figure S2.** Scatterplots of the associations between age (x-axis) and total SPQ (left, *R^2^* < 10^-3^, *P*=0.69, df = 247) and global MT (right, *R^2^* = 0.04, *P*=0.002, df = 247).

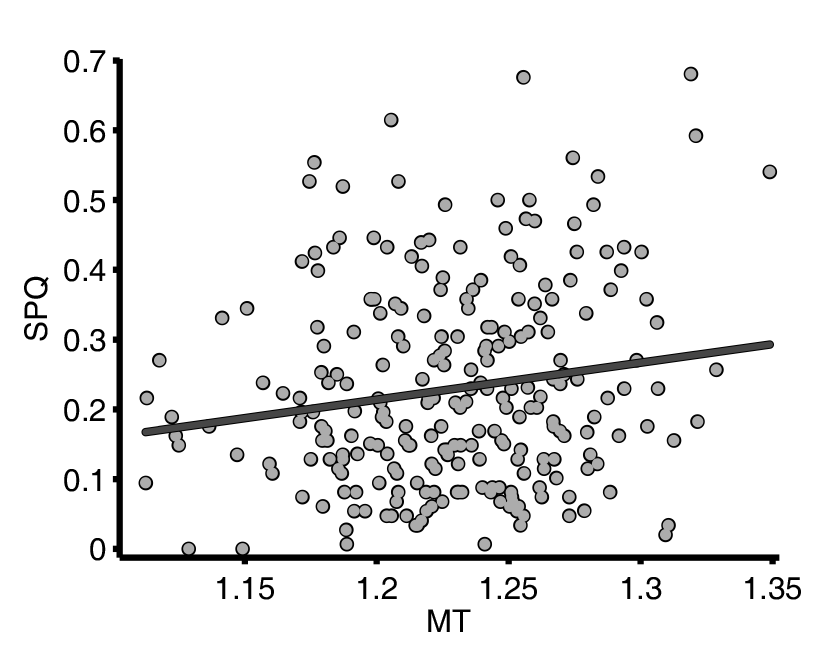

**Figure S3.** Scatterplot of total SPQ score (y-axis) versus global MT (averaged across regions) for 248 participants (*R*^2^ = 0.02; *P* = 0.015; df = 247).

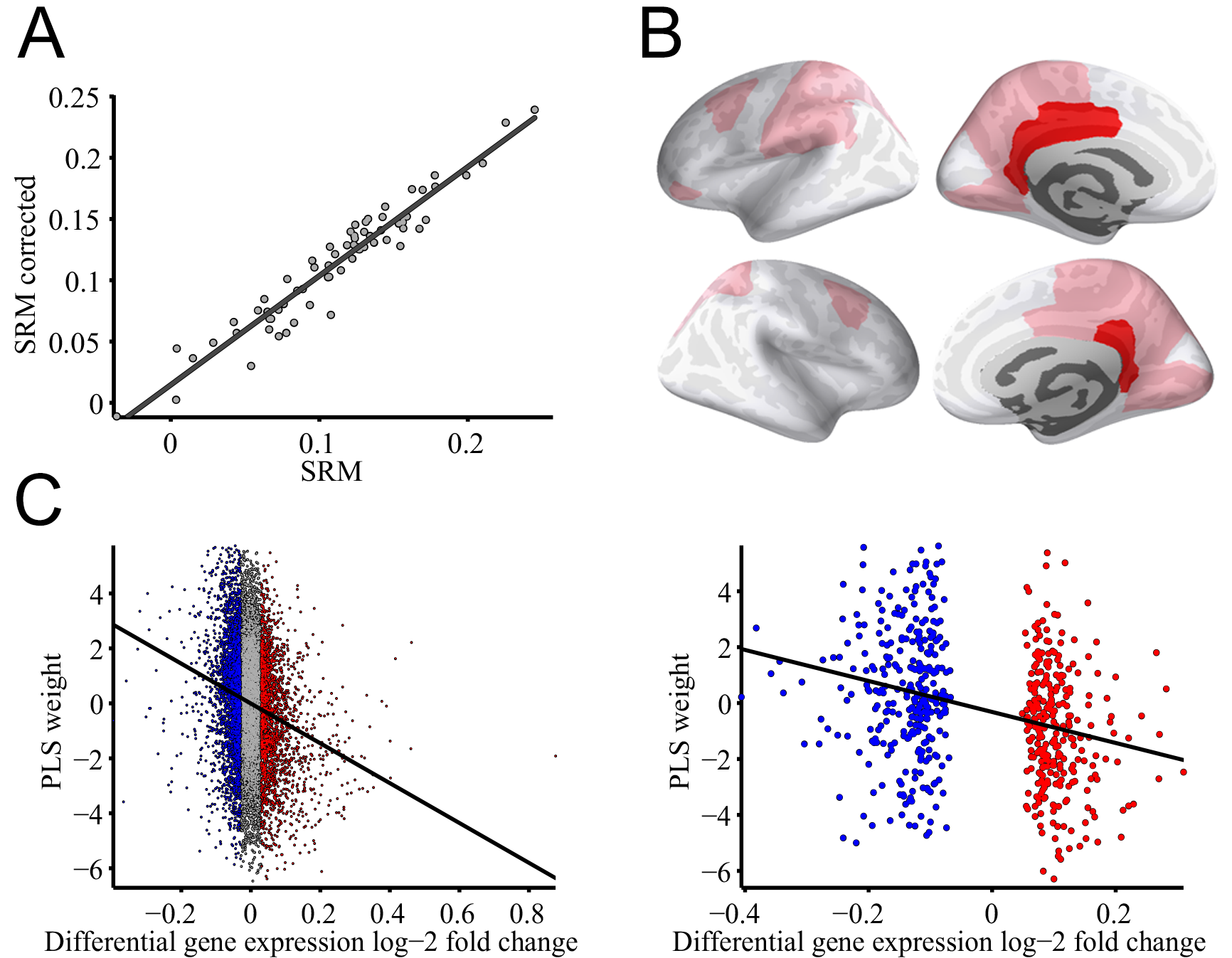

**Figure S4. Schizotypy-related magnetization (SRM) after regressing out the effect of age, gender, scanning site, socio-economic status and total brain volume.** (a) Scatterplot of SRM before and after removing the effect of the covariates (*R^2^* = 0.92, P < 10^-37^, df = 66). (b) Cortical surface maps highlighting areas where total SPQ score was significantly positively correlated with regional MT after controlling for age, gender, scanning site, socio-economic status and total volume by regression: pink regions had nominally significant schizotypy-related magnetization (SRM) (two-tailed *P* < 0.05); red regions had significant SRM controlled for multiple comparisons over 68 cortical regions tested (FDR < 0.05). (c) The weight of each gene on the first PLS component calculated from SRM after regressing out age, gender and total brain volume was significantly negatively correlated with differential gene expression post mortem in schizophrenia according to prior data reported by Gandal et al (2018) (Spearman’s rank correlation, *ρ* = -0.16, Bonferroni-corrected P_adj_ < 10^-61^, df = 11111) and by Fromer et al (2016) (Spearman’s rank correlation, *ρ* = -0.29, P_adj_ < 10^-12^, df = 586; for this dataset only significantly different expression values have been reported (10)).

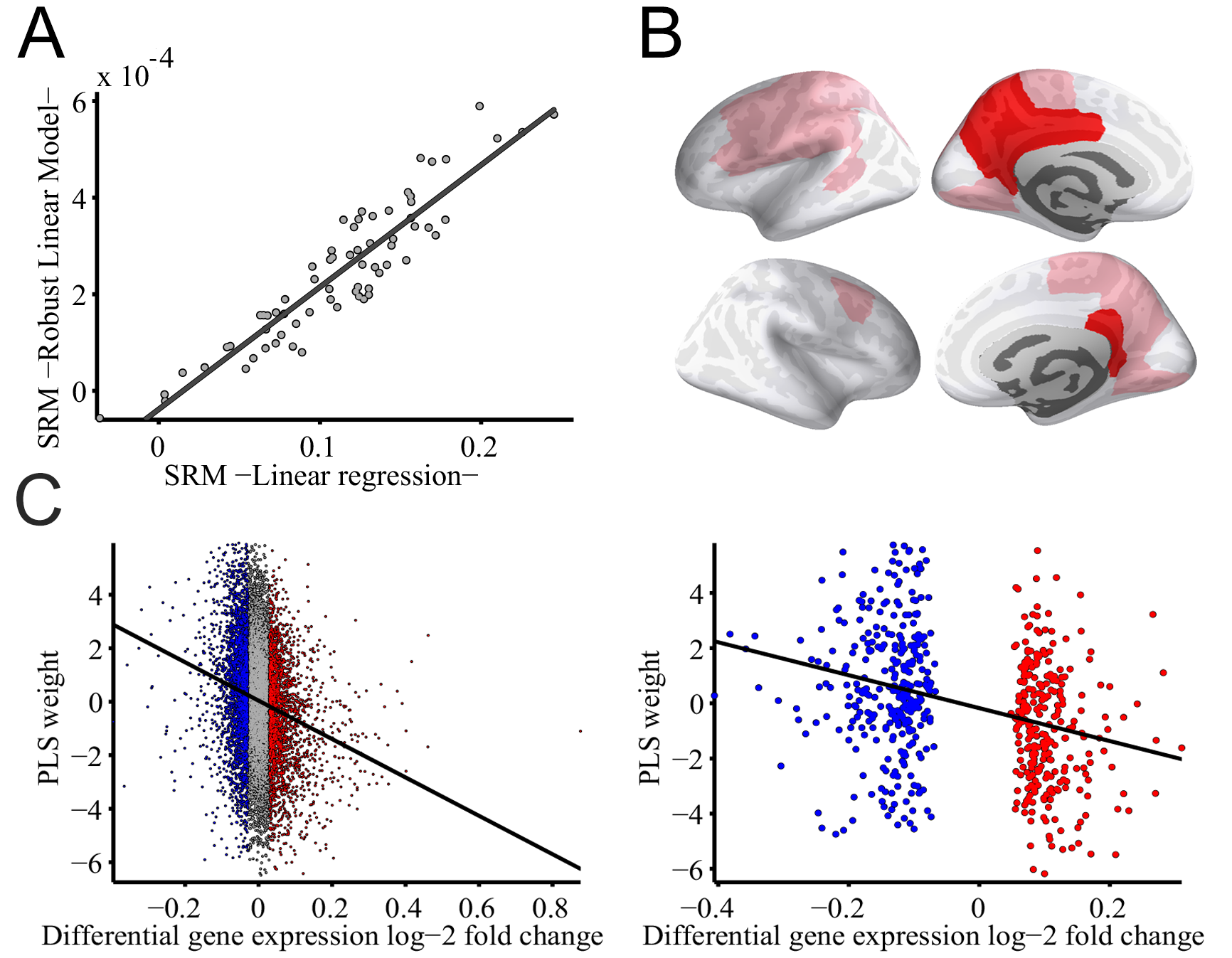

**Figure S5. Schizotypy-related magnetization (SRM) calculated using robust linear models.** (a) Scatterplot of SRM calculated using linear regression and robust linear models (*R^2^* = 0.85, P < 10^-28^, df = 66). (b) Cortical surface maps highlighting areas where total SPQ score was significantly positively correlated with regional MT according to the robust linear regression: pink regions had nominally significant schizotypy-related magnetization (SRM) (two-tailed *P* < 0.05); red regions had significant SRM controlled for multiple comparisons over 68 cortical regions tested (FDR < 0.05). (c) The weight of each gene on the first PLS component calculated from SRM was significantly negatively correlated with differential gene expression post mortem in schizophrenia according to prior data reported by Gandal et al (2018) (Spearman’s rank correlation, *ρ* = -0.18, Bonferroni-corrected P_adj_ < 10^-80^, df = 11111) and by Fromer et al (2016) (Spearman’s rank correlation, *ρ* = -0.28, P_adj_ < 10^-11^, df = 586; for this dataset only significantly different expression values have been reported (10)).

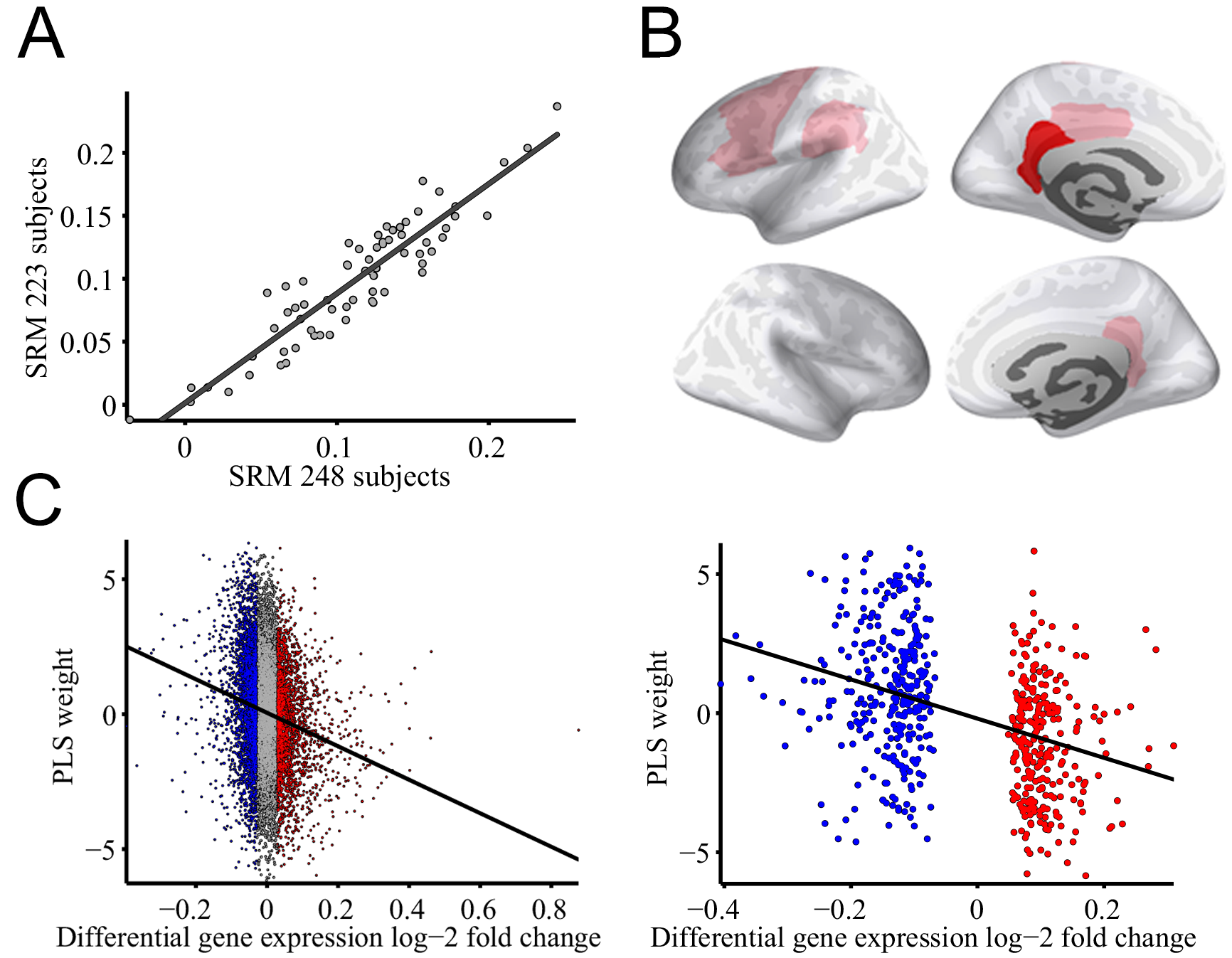

**Figure S6. Schizotypy-related magnetization (SRM) calculated after removing the top 10% of the subject with worst image quality.** (a) Scatterplot of SRM calculated with the original cohort 248 participant and SRM computed after removing those 25 images with highest Euler index (*R^2^* = 0.85, P < 10^-28^, df = 66). (b) Cortical surface maps highlighting areas where total SPQ score calculated using 248 participants was significantly positively correlated with regional MT: pink regions had nominally significant schizotypy-related magnetization (SRM) (two-tailed *P* < 0.05); red regions had significant SRM controlled for multiple comparisons over 68 cortical regions tested (FDR < 0.05). (c) The weight of each gene on the first PLS component calculated from SRM was significantly negatively correlated with differential gene expression post mortem in schizophrenia according to prior data reported by Gandal et al (2018) (Spearman’s rank correlation, *ρ* = -0.16, Bonferroni-corrected P_adj_ < 10^-62^, df = 11111) and by Fromer et al (2016) (Spearman’s rank correlation, *ρ* = -0.36, P_adj_ < 10^-20^, df = 586; for this dataset only significantly different expression values have been reported (10)).

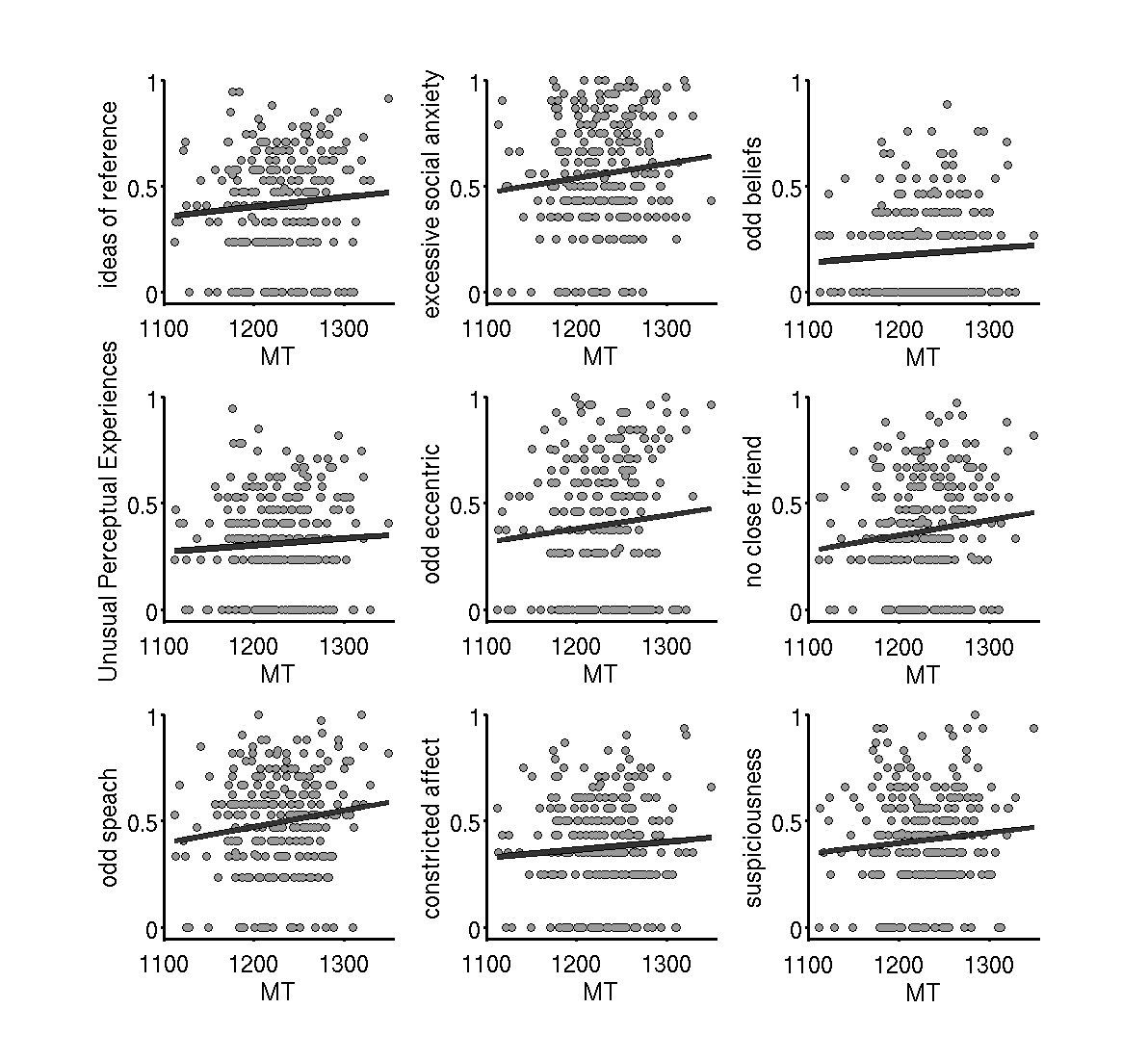

**Figure S7. Association between global MT (averaged across all regions) and SPQ subscales for 248 participants.** None of the correlations were significant after FDR correction (ideas of reference *R^2^*=0.013, *P*=0.17; excessive social anxiety *R^2^*=0.011, *P*=0.19; odd beliefs *R^2^*=0.008, *P*=0.24; unusual perceptual experiences *R^2^*=0.005, *P*=0.28; odd eccentric *R^2^*=0.023, *P*=0.06; no close friend *R^2^*=0.024, *P*=0.06; odd speech *R^2^*=0.022, *P*=0.06; constricted affect *R^2^*=0.005, *P*=0.27; suspiciousness *R^2^*=0.005, *P*=0.27; df = 247).

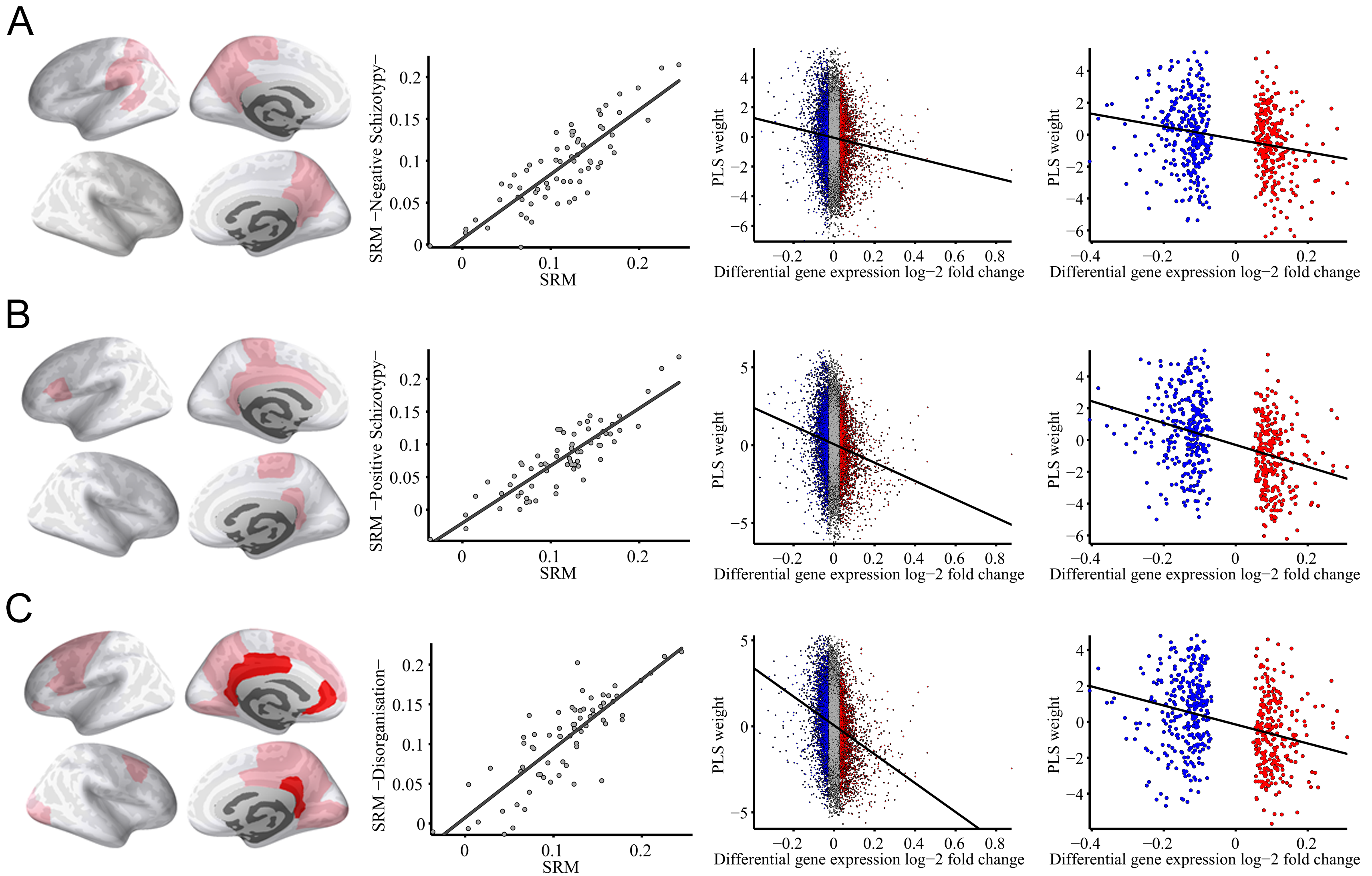

**Figure S8. Association between MT and the three factor models of schizotypy:** **(a) Negative Schizotypy factor, (b) Positive Schizotypy factor and (c) Disorganisation.** Cortical surface maps highlighting areas where each factor score was significantly positively correlated with regional MT (pink, two-tailed *P* < 0.05; red, FDR < 0.05; First column). Scatterplot of SRM calculated from total SPQ compared with SRM based on each individual factor (*R^2^* = 0.70, P < 10^-18^ for Negative Schizotypy, *R^2^* = 0.77, P < 10^-22^ for Positive Schizotypy, *R^2^* = 0.69, P < 10^-17^ for Disorganisation; df=67; Second column). The weight of each gene on the first PLS calculated from SRM based on each individual factor was correlated with differential gene expression post mortem in schizophrenia (Negative Schizotypy: *ρ* = -0.06, P_adj_ < 10^-10^ and *ρ* = -0.24, P_adj_ < 10^-8^ for Gandal -third column- and Fromer -fourth column- differential expression respectivetly; Positive Schizotypy: *ρ* = -0.13, P_adj_ < 10^-47^ and *ρ* = -0.36, P_adj_ < 10^-18^; Disorganisation: *ρ* = -0.22, P_adj_ < 10^-121^ and *ρ* = -0.28, P_adj_ < 10^-12^).

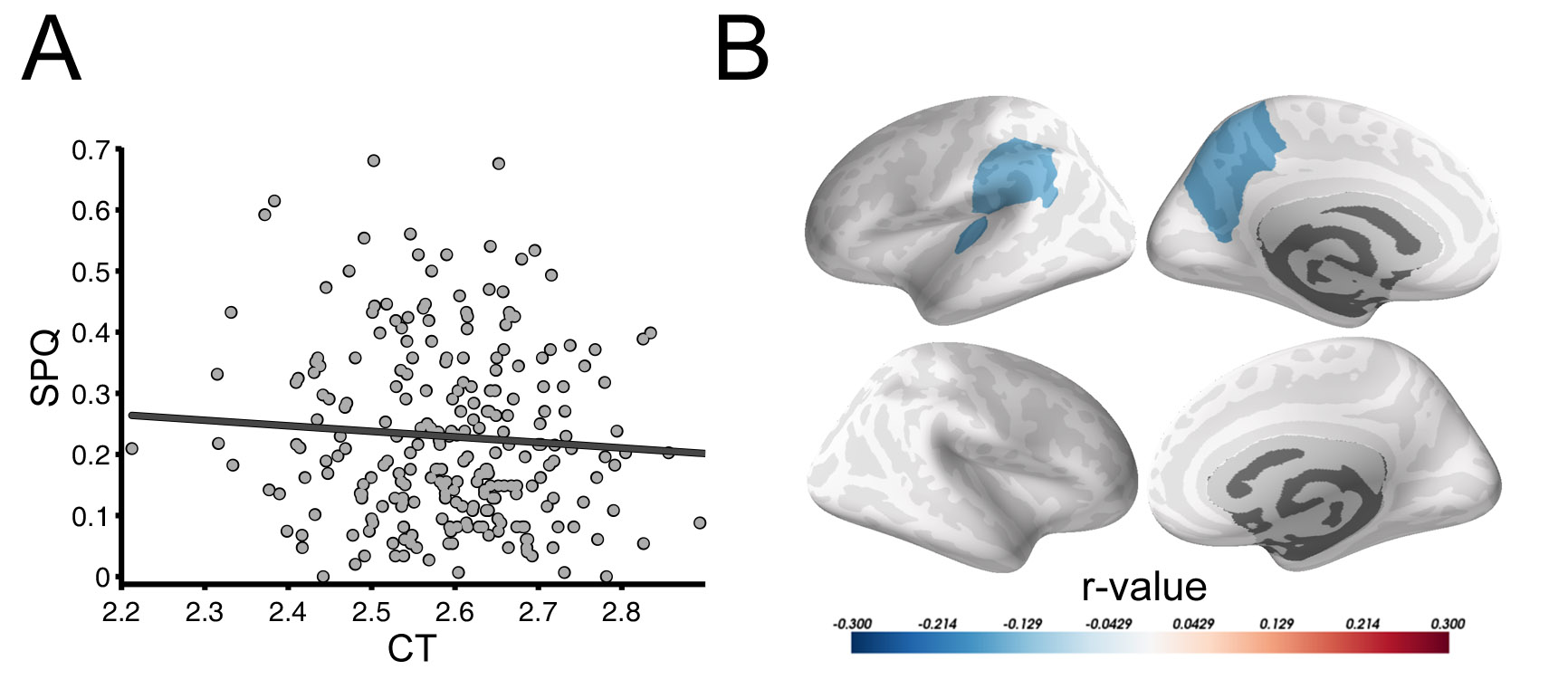

**Figure S9. Schizotypy-dependent cortical thickness of parietal lobe.** (a) Scatterplot of SPQ total score versus global mean CT (averaged over all regions; *R*^2^ = 0.005; *P* = 0.27; df = 247). (b) Cortical surface maps highlighting cortical areas where total SPQ score was significantly negatively correlated with regional CT: blue regions had nominally significant SPQ/CT correlation (*P*<0.025, uncorrected); however, none of these regional associations survived correction for multiple comparisons.

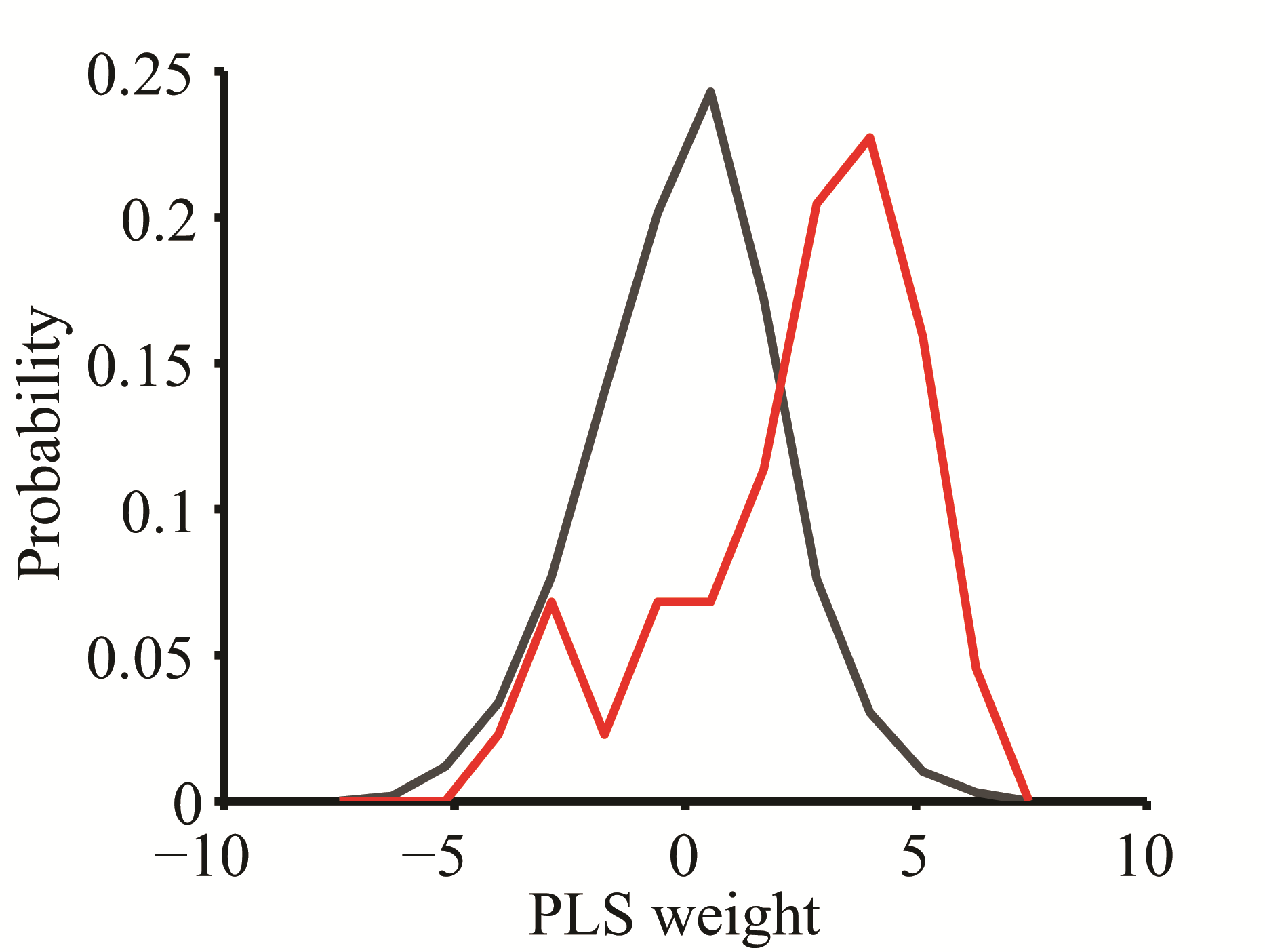

**Figure S10. Distribution of PLS weights calculated from SRM for all genes (black) and genes that correlate with macroscale disconnectivity in schizophrenia as reported by (11).** The 156 genes showing the strongest correlation with white-matter connectivity changes in schizophrenia described by (11) had significantly higher PLS weight that the rest of genes (t-test, *P* < 10^-14^) and were more frequently ranked at the top of the PLS component than expected by chance (permutation test, *P* < 10^-4^).

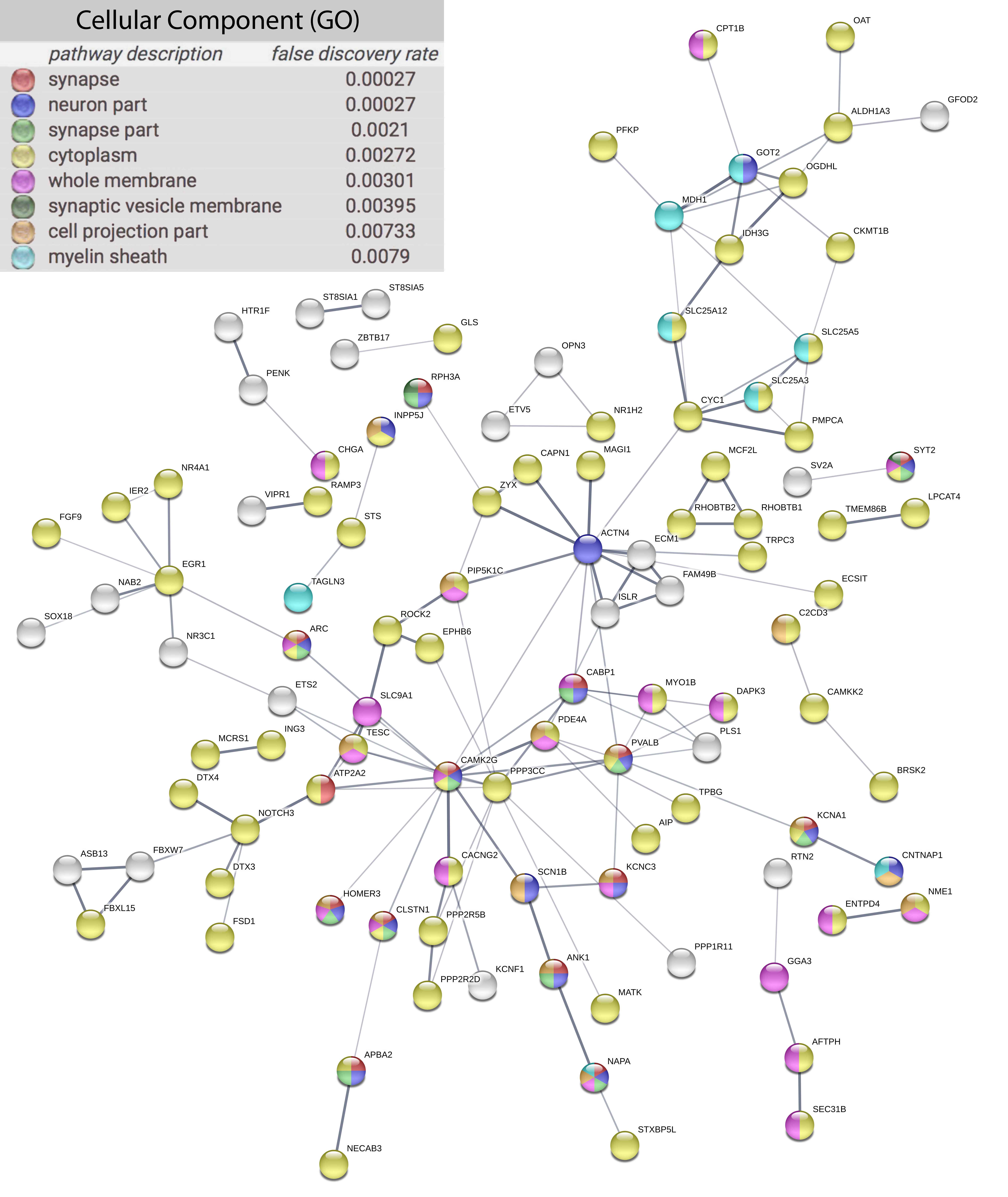

**Figure S11. Protein-protein interaction network for the set of 213 proteins coded by genes associated with both schizotypy-related magnetization and post-mortem brain transcription in schizophrenia.** Nodes represent genes that were both (i) down-regulated in brain tissue from 159 patients with schizophrenia (12); and (ii) positively weighted on the PLS component most strongly associated with schizotypy-related myelination in 248 healthy adolescents. Edges represent known protein-protein interactions and weights are proportional to the STRING confidence score > 0.4 (13). Proteins are coloured by their participation in the most highly enriched GO cellular component terms; all *P*-values are reported after FDR correction. See <https://version-10-5.string-db.org/cgi/network.pl?taskId=RMpA04wbWG8k> for a full interactive map of PPI network.

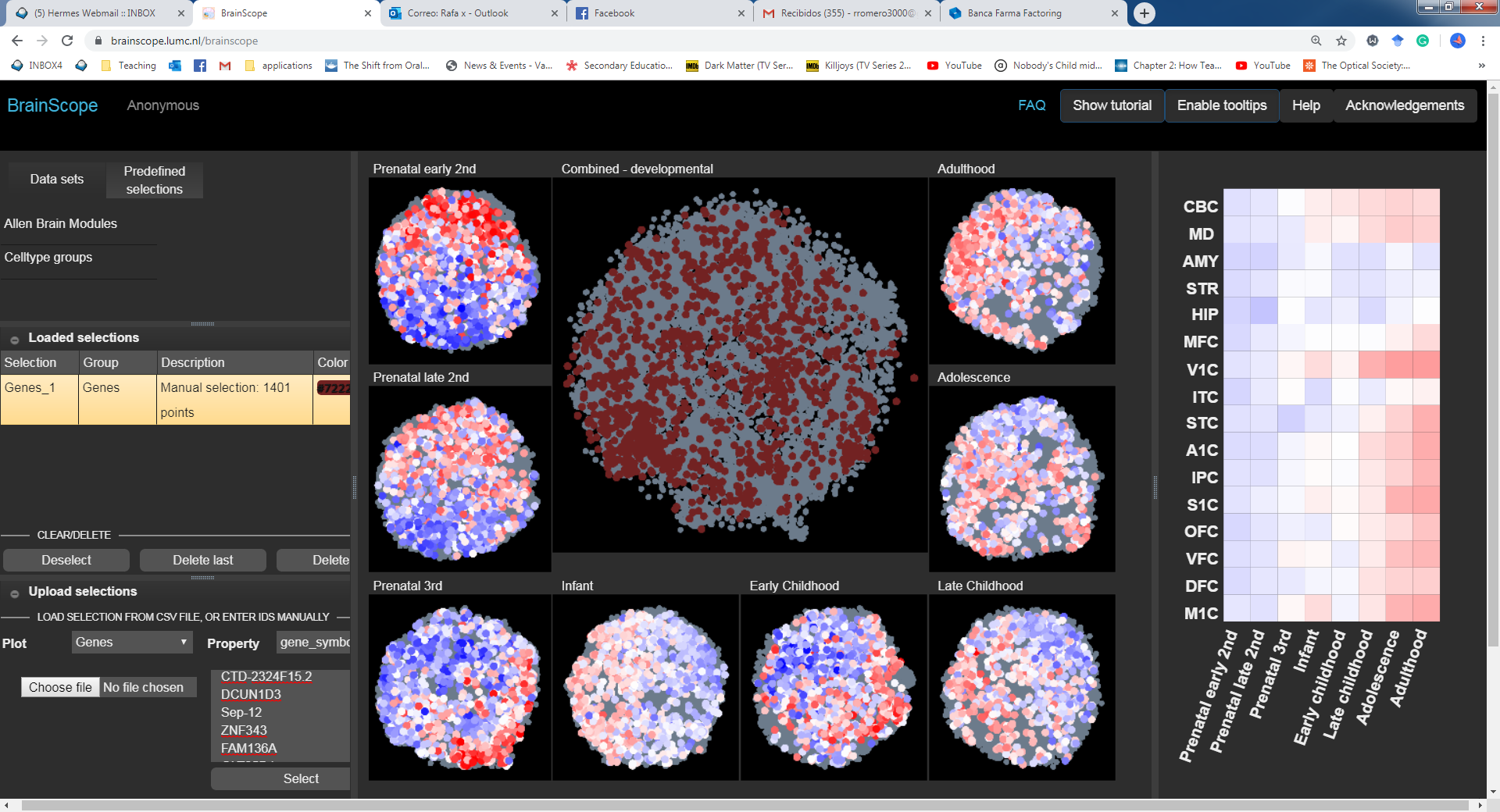

**Figure S12. Expression changes of positively-weighted SRM genes across developmental stages.** Average gene expression values of positively-weighted SRM calculated form the Developing Human Transcriptome included in the BrainSpan Atlas (<http://brainspan.org./static/home>). CBC, cerebellar cortex; MD, mediodorsal nucleus of the thalamus; AMY, amygdala; STR, striatum; HIP, hippocampus; MFC, medial prefrontal cortex; V1C, primary visual (V1) cortex; ITC, inferior temporal cortex; STC, superior temporal cortex; A1C, primary auditory (A1) cortex; IPC, posterior inferior parietal cortex; S1C, primary somatosensory (S1) cortex; OFC, orbital prefrontal cortex; VFC, ventrolateral prefrontal cortex; DFC, dorsolateral prefrontal cortex; M1C, primary motor (M1) cortex. Figure generated using BrainScope (<https://brainscope.lumc.nl/brainscope>).

### Supplementary Tables

| **Age (yrs)** | **Gender** | **N** | **IR** | **ESA** | **OBMT** | **UPE** | **OEB** | **NCF** | **OS** | **CA** | **S** | **Total SPQ** |
| --- | --- | --- | --- | --- | --- | --- | --- | --- | --- | --- | --- | --- |
| 14 to 15 | M | 27 | 0.14 | 0.21 | 0.02 | 0.10 | 0.18 | 0.07 | 0.18 | 0.07 | 0.18 | **0.09** |
|  | F | 25 | 0.22 | 0.32 | 0.02 | 0.05 | 0.17 | 0.07 | 0.19 | 0.05 | 0.12 | **0.08** |
| 16 to 17 | M | 25 | 0.14 | 0.23 | 0.06 | 0.09 | 0.26 | 0.12 | 0.20 | 0.11 | 0.14 | **0.10** |
|  | F | 26 | 0.17 | 0.29 | 0.08 | 0.07 | 0.13 | 0.07 | 0.16 | 0.06 | 0.13 | **0.09** |
| 18 to 19 | M | 25 | 0.18 | 0.26 | 0.03 | 0.10 | 0.16 | 0.11 | 0.20 | 0.14 | 0.23 | **0.12** |
|  | F | 27 | 0.15 | 0.30 | 0.04 | 0.06 | 0.12 | 0.11 | 0.13 | 0.06 | 0.15 | **0.08** |
| 20 to 21 | M | 21 | 0.10 | 0.27 | 0.05 | 0.12 | 0.33 | 0.06 | 0.21 | 0.10 | 0.08 | **0.10** |
|  | F | 22 | 0.11 | 0.20 | 0.04 | 0.07 | 0.13 | 0.04 | 0.14 | 0.04 | 0.16 | **0.07** |
| 22 to 25 | M | 27 | 0.10 | 0.20 | 0.03 | 0.06 | 0.19 | 0.17 | 0.20 | 0.12 | 0.12 | **0.09** |
|  | F | 23 | 0.21 | 0.40 | 0.06 | 0.06 | 0.25 | 0.13 | 0.21 | 0.12 | 0.17 | **0.12** |

**Table S1: Schizotypal Personality Questionnaire: total score and subscale scores stratified by age and sex**. IR = Ideas of References; ESA = Excessive Social Anxiety; OBMT = Odd Beliefs or Magical Thinking; UPE = Unusual Perceptual Experiences; OEB = Odd or Eccentric Behavior; NCF = No Close Friends; OS = Odd Speech; CA = Constricted Affect; S = Suspiciousness.

|  | Number of genes |
| --- | --- |
| Astrocyte | 280 |
| Endothelial | 177 |
| GabaPV | 15 |
| GabaRelnCalb | 15 |
| GabaVIPReln | 33 |
| Layer 2 3 Pyra | 4 |
| Layer 4 Pyra | 5 |
| Layer 6a Pyra | 6 |
| Layer 6b Pyra | 9 |
| Microglia | 284 |
| Microglia_activation | 118 |
| Microglia_deactivation | 199 |
| Oligo | 152 |
| OligoPrecursors | 185 |
| Pyramidal | 35 |
| PyramidalCorticoThalam | 2 |
| Pyramidal_Glt_25d2 | 3 |
| Pyramidal_S100a10 | 2 |

**Table S2.** Cell-type expression profiles defined by (10). Each cell-type was associated with the differential expression of a subset of genes.

|  | **Gandal Up-reg** | **Gandal Down-reg** | **Fromer Up-reg** | **Fromer Down-reg** |
| --- | --- | --- | --- | --- |
| **Gandal Up-reg** | 845 | 0 | 39 | 11 |
| **Gandal Down-reg** |  | 1177 | 4 | 76 |
| **Fromer Up-reg** |  |  | 304 | 0 |
| **Fromer Down-reg** |  |  |  | 345 |

**Table S3.** Number of schizophrenia-related genes overlapping between lists of up and down-regulated genes defined by (12) and up and down-regulated genes defined by (14).

| **Region** | ***r*-value** | ***p*-value** |
| --- | --- | --- |
| LH isthmuscingulate | 0.24506 | 9.65E-05 |
| RH isthmuscingulate | 0.22557 | 0.000343 |
| LH posteriorcingulate | 0.21008 | 0.000872 |
| LH precuneus | 0.19896 | 0.001639 |
| LH supramarginal | 0.17812 | 0.004901 |
| LH paracentral | 0.1778 | 0.004981 |
| RH caudalmiddlefrontal | 0.17176 | 0.0067 |
| RH lingual | 0.1696 | 0.007432 |
| LH precentral | 0.16741 | 0.00825 |
| RH precuneus | 0.16242 | 0.01041 |
| LH postcentral | 0.15889 | 0.01223 |
| LH parsopercularis | 0.15656 | 0.013577 |
| LH superiorparietal | 0.15644 | 0.01365 |
| LH lingual | 0.15627 | 0.013755 |
| LH bankssts | 0.15461 | 0.014801 |
| LH caudalmiddlefrontal | 0.15349 | 0.015552 |
| RH precentral | 0.14538 | 0.022019 |
| RH paracentral | 0.14435 | 0.022985 |
| RH posteriorcingulate | 0.14266 | 0.024656 |
| LH caudalanteriorcingulate | 0.14153 | 0.025827 |
| RH medialorbitofrontal | 0.13686 | 0.031203 |
| LH superiorfrontal | 0.13413 | 0.03476 |
| RH pericalcarine | 0.13276 | 0.036671 |
| RH superiorparietal | 0.1312 | 0.038951 |
| LH lateralorbitofrontal | 0.13029 | 0.04034 |
| RH lateralorbitofrontal | 0.13029 | 0.040341 |
| LH rostralanteriorcingulate | 0.12723 | 0.045321 |
| RH parstriangularis | 0.12629 | 0.046953 |
| LH inferiorparietal | 0.12592 | 0.047608 |
| LH parsorbitalis | 0.12421 | 0.050721 |
| RH supramarginal | 0.12389 | 0.05134 |
| LH rostralmiddlefrontal | 0.12364 | 0.051801 |
| LH transversetemporal | 0.12332 | 0.052415 |
| RH superiorfrontal | 0.12224 | 0.054534 |
| RH lateraloccipital | 0.12104 | 0.056973 |
| RH postcentral | 0.11864 | 0.0621 |
| RH insula | 0.11452 | 0.071826 |
| RH parsorbitalis | 0.1107 | 0.081872 |
| LH insula | 0.10785 | 0.090124 |
| LH pericalcarine | 0.10716 | 0.092208 |
| LH parstriangularis | 0.1067 | 0.093625 |
| LH middletemporal | 0.10632 | 0.094807 |
| LH cuneus | 0.1058 | 0.096429 |
| RH fusiform | 0.096602 | 0.12923 |
| RH inferiorparietal | 0.095275 | 0.1346 |
| RH rostralmiddlefrontal | 0.093595 | 0.14164 |
| LH frontalpole | 0.088892 | 0.16285 |
| LH parahippocampal | 0.085186 | 0.18117 |
| RH entorhinal | 0.083059 | 0.19235 |
| LH lateraloccipital | 0.078353 | 0.21886 |
| RH parsopercularis | 0.077761 | 0.22237 |
| LH medialorbitofrontal | 0.076083 | 0.23254 |
| LH superiortemporal | 0.072809 | 0.25332 |
| RH rostralanteriorcingulate | 0.072621 | 0.25455 |
| RH bankssts | 0.067464 | 0.28994 |
| RH cuneus | 0.066513 | 0.29681 |
| RH caudalanteriorcingulate | 0.066168 | 0.29932 |
| LH inferiortemporal | 0.065014 | 0.30784 |
| RH inferiortemporal | 0.062987 | 0.32321 |
| RH temporalpole | 0.058726 | 0.35708 |
| RH frontalpole | 0.054053 | 0.39669 |
| RH transversetemporal | 0.044336 | 0.48704 |
| LH fusiform | 0.042373 | 0.50656 |
| LH entorhinal | -0.03653 | 0.56697 |
| RH parahippocampal | 0.028528 | 0.65481 |
| RH middletemporal | 0.014899 | 0.8154 |
| LH temporalpole | 0.004035 | 0.94959 |
| RH superiortemporal | 0.003549 | 0.95566 |

**Table S4.** Correlation between total SPQ and regional MT. Red rows highlight significant correlations (*P*<0.05, FDR corrected).

| **Gene Name** | **Schizophrenia-related list** | **PLS1**  ***Z*-score** | **PLS1**  **Adjusted *P*-value** | **PLS1**  **Ranking out of 20,647** | **Inter-regional**  **correlation** | **Inter-regional**  ***P*-value** |
| --- | --- | --- | --- | --- | --- | --- |
| *IER2* | GDR | 6.41 | 1.00E-06 | 2 | 0.43 | 2.26E-02 |
| *ESRRA* | GDR | 6.12 | 1.00E-06 | 9 | 0.50 | 7.91E-03 |
| *CAMK2G* | GDR | 6.03 | 1.00E-06 | 22 | 0.47 | 1.18E-02 |
| *PDE4A* | GDR | 6.03 | 1.00E-06 | 23 | 0.44 | 1.94E-02 |
| *TRPC3* | GDR | 5.99 | 1.00E-06 | 25 | 0.36 | 6.82E-02 |
| *ESRRG* | GDR | 5.99 | 1.00E-06 | 26 | 0.43 | 2.23E-02 |
| *ANK1* | GDR | 5.96 | 1.00E-06 | 27 | 0.45 | 1.76E-02 |
| *ECM1* | GDR | 5.94 | 1.00E-06 | 28 | 0.45 | 1.82E-02 |
| *SCN1B* | GDR | 5.92 | 1.00E-06 | 29 | 0.40 | 3.81E-02 |
| *KCNA1* | GDR | 5.76 | 2.00E-06 | 38 | 0.45 | 1.74E-02 |
| *SLC25A5* | GDR | 5.74 | 2.00E-06 | 41 | 0.33 | 9.82E-02 |
| *ING3* | GDR | 5.71 | 3.00E-06 | 45 | 0.37 | 6.02E-02 |
| *KCNC3* | GDR | 5.68 | 3.00E-06 | 48 | 0.42 | 2.92E-02 |
| *SEMA7A* | GDR | 5.63 | 4.00E-06 | 54 | 0.38 | 4.90E-02 |
| *FAM131B* | GDR | 5.51 | 5.00E-06 | 75 | 0.49 | 9.25E-03 |
| *SLC38A1* | GDR | 5.45 | 6.00E-06 | 82 | 0.45 | 1.74E-02 |
| *RHOBTB2* | GDR | 5.39 | 8.00E-06 | 90 | 0.45 | 1.88E-02 |
| *GLS2* | GDR | 5.38 | 8.00E-06 | 91 | 0.38 | 5.34E-02 |
| *RFX5* | GDR | 5.37 | 9.00E-06 | 94 | 0.43 | 2.60E-02 |
| *HTR1F* | GDR | 5.33 | 1.00E-05 | 99 | 0.43 | 2.48E-02 |
| *SPHK2* | GDR | 5.32 | 1.00E-05 | 101 | 0.48 | 9.25E-03 |
| *ETS2* | GDR | 5.28 | 1.30E-05 | 106 | 0.34 | 8.44E-02 |
| *SYT2* | GDR | 5.16 | 2.10E-05 | 116 | 0.38 | 4.80E-02 |
| *CRABP1* | GDR | 5.13 | 2.40E-05 | 121 | 0.34 | 9.52E-02 |
| *DNAJC4* | GDR | 5.12 | 2.60E-05 | 122 | 0.36 | 6.68E-02 |
| *CNTNAP1* | GDR | 5.11 | 2.70E-05 | 126 | 0.43 | 2.62E-02 |
| *SCRT1* | GDR | 5.07 | 3.10E-05 | 132 | 0.42 | 2.89E-02 |
| *OGDHL* | GDR | 5.00 | 4.00E-05 | 143 | 0.36 | 7.19E-02 |
| *FGF9* | GDR | 4.99 | 4.20E-05 | 150 | 0.40 | 3.97E-02 |
| *TRAF3* | GDR | 4.97 | 4.30E-05 | 155 | 0.48 | 9.95E-03 |
| *DCUN1D2* | GDR | 4.94 | 5.00E-05 | 163 | 0.40 | 3.97E-02 |
| *ST8SIA5* | GDR | 4.90 | 5.80E-05 | 167 | 0.37 | 6.02E-02 |
| *HECA* | GDR | 4.90 | 6.00E-05 | 169 | 0.40 | 3.63E-02 |
| *GFRA2* | GDR | 4.86 | 7.00E-05 | 173 | 0.43 | 2.23E-02 |
| *SLC7A8* | GDR | 4.85 | 7.30E-05 | 178 | 0.48 | 9.68E-03 |
| *CKMT1B* | GDR | 4.84 | 7.40E-05 | 180 | 0.37 | 5.92E-02 |
| *OSBPL3* | GDR | 4.82 | 8.00E-05 | 183 | 0.45 | 1.82E-02 |
| *SYT12* | GDR | 4.76 | 1.03E-04 | 199 | 0.41 | 3.15E-02 |
| *CABP1* | GDR | 4.74 | 1.08E-04 | 207 | 0.39 | 4.40E-02 |
| *LUZP1* | GDR | 4.74 | 1.08E-04 | 208 | 0.36 | 7.23E-02 |
| *PLXDC1* | GDR | 4.73 | 1.09E-04 | 210 | 0.41 | 3.19E-02 |
| *RAB11FIP5* | GDR | 4.66 | 1.41E-04 | 237 | 0.34 | 9.60E-02 |
| *ASB13* | GDR | 4.65 | 1.44E-04 | 239 | 0.36 | 6.51E-02 |
| *SLC25A12* | GDR | 4.63 | 1.55E-04 | 240 | 0.33 | 9.82E-02 |
| *DDHD2* | GDR | 4.63 | 1.59E-04 | 241 | 0.42 | 2.66E-02 |
| *ZMAT4* | GDR | 4.61 | 1.68E-04 | 248 | 0.37 | 6.27E-02 |
| *MAGI1* | GDR | 4.59 | 1.77E-04 | 253 | 0.28 | 1.82E-01 |
| *DPP8* | GDR | 4.58 | 1.84E-04 | 256 | 0.35 | 8.41E-02 |
| *PRDM1* | GDR | 4.58 | 1.86E-04 | 260 | 0.45 | 1.81E-02 |
| *SV2A* | GDR | 4.58 | 1.87E-04 | 261 | 0.37 | 5.68E-02 |
| *SLC20A1* | GDR | 4.55 | 2.07E-04 | 269 | 0.46 | 1.44E-02 |
| *ARC* | GDR | 4.55 | 2.09E-04 | 270 | 0.40 | 3.92E-02 |
| *OSBP2* | GDR | 4.54 | 2.10E-04 | 271 | 0.37 | 6.24E-02 |
| *FBXW7* | GDR | 4.49 | 2.55E-04 | 282 | 0.38 | 5.36E-02 |
| *CPT1B* | GDR | 4.49 | 2.64E-04 | 285 | 0.37 | 5.68E-02 |
| *CHRD* | GDR | 4.45 | 3.03E-04 | 295 | 0.32 | 1.12E-01 |
| *SV2C* | GDR | 4.43 | 3.26E-04 | 299 | 0.34 | 8.44E-02 |
| *NR3C1* | GDR | 4.40 | 3.58E-04 | 315 | 0.40 | 3.55E-02 |
| *CAMKK2* | GDR | 4.37 | 3.90E-04 | 323 | 0.39 | 4.57E-02 |
| *DEPDC5* | GDR | 4.37 | 3.91E-04 | 324 | 0.37 | 5.71E-02 |
| *ZYX* | GDR | 4.35 | 4.23E-04 | 335 | 0.38 | 4.97E-02 |
| *ATG16L1* | GDR | 4.35 | 4.24E-04 | 336 | 0.29 | 1.73E-01 |
| *HHAT* | GDR | 4.33 | 4.51E-04 | 344 | 0.55 | 4.73E-03 |
| *ROCK2* | GDR | 4.33 | 4.54E-04 | 345 | 0.34 | 8.75E-02 |
| *PNMA3* | GDR | 4.31 | 4.70E-04 | 356 | 0.33 | 1.03E-01 |
| *ST8SIA1* | GDR | 4.31 | 4.70E-04 | 358 | 0.31 | 1.25E-01 |
| *POU6F1* | GDR | 4.30 | 4.85E-04 | 365 | 0.31 | 1.28E-01 |
| *BRSK2* | GDR | 4.28 | 5.16E-04 | 374 | 0.34 | 8.43E-02 |
| *LPCAT4* | GDR | 4.27 | 5.30E-04 | 380 | 0.32 | 1.23E-01 |
| *HR* | GDR | 4.27 | 5.34E-04 | 382 | 0.37 | 5.46E-02 |
| *NARS* | GDR | 4.24 | 5.80E-04 | 398 | 0.33 | 1.05E-01 |
| *ATRNL1* | GDR | 4.22 | 6.26E-04 | 406 | 0.33 | 1.04E-01 |
| *NUAK1* | GDR | 4.22 | 6.26E-04 | 407 | 0.33 | 9.92E-02 |
| *ALDH1A3* | GDR | 4.22 | 6.28E-04 | 410 | 0.29 | 1.73E-01 |
| *PPP3CC* | GDR | 4.15 | 7.93E-04 | 440 | 0.25 | 2.67E-01 |
| *ISLR* | GDR | 4.13 | 8.28E-04 | 449 | 0.38 | 5.08E-02 |
| *TMEM127* | GDR | 4.07 | 9.98E-04 | 482 | 0.27 | 2.24E-01 |
| *C6orf106* | GDR | 4.02 | 1.19E-03 | 501 | 0.33 | 9.73E-02 |
| *PFKP* | GDR | 4.01 | 1.22E-03 | 508 | 0.32 | 1.14E-01 |
| *FNDC4* | GDR | 4.01 | 1.23E-03 | 509 | 0.30 | 1.53E-01 |
| *PVALB* | GDR | 4.01 | 1.25E-03 | 511 | 0.31 | 1.36E-01 |
| *RUNDC3B* | GDR | 3.99 | 1.30E-03 | 520 | 0.25 | 2.61E-01 |
| *ZNF385D* | GDR | 3.98 | 1.36E-03 | 529 | 0.36 | 6.93E-02 |
| *RHOBTB1* | GDR | 3.97 | 1.40E-03 | 533 | 0.36 | 7.10E-02 |
| *EPHB6* | GDR | 3.93 | 1.59E-03 | 548 | 0.42 | 2.92E-02 |
| *LGI2* | GDR | 3.93 | 1.60E-03 | 556 | 0.33 | 1.07E-01 |
| *LAPTM4B* | GDR | 3.92 | 1.65E-03 | 559 | 0.30 | 1.49E-01 |
| *GOT2* | GDR | 3.88 | 1.85E-03 | 579 | 0.27 | 2.02E-01 |
| *ANKS1A* | GDR | 3.85 | 2.05E-03 | 597 | 0.27 | 2.14E-01 |
| *SCAMP5* | GDR | 3.84 | 2.12E-03 | 605 | 0.32 | 1.18E-01 |
| *IDH3G* | GDR | 3.83 | 2.16E-03 | 608 | 0.31 | 1.26E-01 |
| *STAU2* | GDR | 3.82 | 2.22E-03 | 624 | 0.37 | 6.13E-02 |
| *ANXA6* | GDR | 3.82 | 2.22E-03 | 626 | 0.25 | 2.62E-01 |
| *SMYD2* | GDR | 3.79 | 2.44E-03 | 646 | 0.33 | 1.04E-01 |
| *PITPNA* | GDR | 3.77 | 2.58E-03 | 659 | 0.29 | 1.70E-01 |
| *STS* | GDR | 3.74 | 2.76E-03 | 679 | 0.23 | 3.02E-01 |
| *RAMP3* | GDR | 3.74 | 2.76E-03 | 681 | 0.42 | 3.10E-02 |
| *PCGF1* | GDR | 3.73 | 2.83E-03 | 688 | 0.33 | 1.03E-01 |
| *VIPR1* | GDR | 3.69 | 3.30E-03 | 716 | 0.30 | 1.58E-01 |
| *RTN2* | GDR | 3.68 | 3.35E-03 | 722 | 0.39 | 4.38E-02 |
| *TPBG* | GDR | 3.66 | 3.50E-03 | 732 | 0.33 | 1.02E-01 |
| *MED22* | GDR | 3.66 | 3.50E-03 | 733 | 0.24 | 2.85E-01 |
| *MAP4K2* | GDR | 3.66 | 3.51E-03 | 736 | 0.33 | 1.00E-01 |
| *PKNOX2* | GDR | 3.66 | 3.57E-03 | 741 | 0.28 | 1.84E-01 |
| *PHLDA2* | GDR | 3.65 | 3.60E-03 | 742 | 0.30 | 1.58E-01 |
| *HOMER3* | GDR | 3.65 | 3.62E-03 | 744 | 0.25 | 2.59E-01 |
| *GFOD2* | GDR | 3.64 | 3.77E-03 | 747 | 0.32 | 1.10E-01 |
| *DTX3* | GDR | 3.63 | 3.90E-03 | 754 | 0.25 | 2.67E-01 |
| *OAT* | GDR | 3.63 | 3.92E-03 | 755 | 0.32 | 1.17E-01 |
| *NR4A1* | GDR | 3.61 | 4.13E-03 | 770 | 0.30 | 1.54E-01 |
| *NRN1* | GDR | 3.60 | 4.24E-03 | 779 | 0.27 | 2.21E-01 |
| *HS3ST1* | GDR | 3.59 | 4.40E-03 | 788 | 0.22 | 3.35E-01 |
| *OPN3* | GDR | 3.59 | 4.40E-03 | 789 | 0.39 | 4.54E-02 |
| *ICA1* | GDR | 3.56 | 4.78E-03 | 809 | 0.28 | 1.97E-01 |
| *CHGA* | GDR | 3.55 | 4.81E-03 | 812 | 0.33 | 9.91E-02 |
| *GLS* | GDR | 3.49 | 5.75E-03 | 857 | 0.24 | 2.89E-01 |
| *SRA1* | GDR | 3.48 | 5.93E-03 | 863 | 0.25 | 2.62E-01 |
| *CLSTN1* | GDR | 3.48 | 5.99E-03 | 866 | 0.29 | 1.67E-01 |
| *ETV5* | GDR | 3.45 | 6.58E-03 | 883 | 0.33 | 1.08E-01 |
| *BID* | GDR | 3.43 | 7.02E-03 | 899 | 0.17 | 5.08E-01 |
| *NAB2* | GDR | 3.42 | 7.20E-03 | 912 | 0.28 | 1.80E-01 |
| *EXTL2* | GDR | 3.40 | 7.60E-03 | 922 | 0.32 | 1.20E-01 |
| *C16orf45* | GDR | 3.39 | 7.69E-03 | 930 | 0.25 | 2.55E-01 |
| *KCNK12* | GDR | 3.35 | 8.55E-03 | 975 | 0.25 | 2.63E-01 |
| *PLS1* | GDR | 3.33 | 9.04E-03 | 986 | 0.32 | 1.14E-01 |
| *CRTAC1* | GDR | 3.33 | 9.06E-03 | 987 | 0.32 | 1.12E-01 |
| *TAPBP* | GDR | 3.33 | 9.21E-03 | 990 | 0.29 | 1.72E-01 |
| *SERGEF* | GDR | 3.32 | 9.28E-03 | 994 | 0.39 | 4.67E-02 |
| *SLC9A1* | GDR | 3.32 | 9.45E-03 | 1001 | 0.27 | 2.11E-01 |
| *JOSD1* | GDR | 3.31 | 9.49E-03 | 1005 | 0.30 | 1.49E-01 |
| *NECAB3* | GDR | 3.31 | 9.63E-03 | 1016 | 0.21 | 3.58E-01 |
| *ABCC5* | GDR | 3.30 | 9.82E-03 | 1025 | 0.26 | 2.49E-01 |
| *PEPD* | GDR | 3.30 | 9.83E-03 | 1030 | 0.21 | 3.73E-01 |
| *REEP2* | GDR | 3.29 | 9.95E-03 | 1038 | 0.26 | 2.33E-01 |
| *TMEM86B* | GDR | 3.27 | 1.06E-02 | 1049 | 0.29 | 1.77E-01 |
| *SLC25A3* | GDR | 3.26 | 1.09E-02 | 1064 | 0.26 | 2.35E-01 |
| *ZBTB17* | GDR | 3.25 | 1.11E-02 | 1068 | 0.32 | 1.08E-01 |
| *TESC* | GDR | 3.24 | 1.13E-02 | 1076 | 0.27 | 2.09E-01 |
| *C2CD3* | GDR | 3.24 | 1.13E-02 | 1079 | 0.28 | 1.98E-01 |
| *MCF2L* | GDR | 3.24 | 1.13E-02 | 1080 | 0.22 | 3.39E-01 |
| *PIP5K1C* | GDR | 3.24 | 1.15E-02 | 1082 | 0.31 | 1.34E-01 |
| *MAP3K13* | GDR | 3.23 | 1.17E-02 | 1089 | 0.29 | 1.66E-01 |
| *C8orf4* | GDR | 3.23 | 1.18E-02 | 1093 | 0.37 | 6.02E-02 |
| *ACTN4* | GDR | 3.22 | 1.21E-02 | 1103 | 0.27 | 2.18E-01 |
| *EGR1* | GDR | 3.22 | 1.21E-02 | 1106 | 0.27 | 2.03E-01 |
| *PPP2R5B* | GDR | 3.21 | 1.22E-02 | 1115 | 0.29 | 1.73E-01 |
| *INPP5J* | GDR | 3.21 | 1.23E-02 | 1119 | 0.25 | 2.72E-01 |
| *LRRC59* | GDR | 3.21 | 1.23E-02 | 1124 | 0.23 | 3.16E-01 |
| *RUSC2* | GDR | 3.20 | 1.24E-02 | 1125 | 0.30 | 1.42E-01 |
| *UBFD1* | GDR | 3.19 | 1.29E-02 | 1143 | 0.25 | 2.63E-01 |
| *LBH* | GDR | 3.18 | 1.31E-02 | 1153 | 0.33 | 1.02E-01 |
| *DUS1L* | GDR | 3.17 | 1.34E-02 | 1160 | 0.27 | 2.23E-01 |
| *NXPH3* | GDR | 3.17 | 1.35E-02 | 1165 | 0.28 | 1.79E-01 |
| *NOTCH3* | GDR | 3.15 | 1.41E-02 | 1188 | 0.32 | 1.09E-01 |
| *RWDD1* | GDR | 3.15 | 1.42E-02 | 1191 | 0.20 | 4.03E-01 |
| *GABRA4* | GDR | 3.15 | 1.42E-02 | 1193 | 0.29 | 1.66E-01 |
| *ATP2A2* | GDR | 3.14 | 1.45E-02 | 1199 | 0.23 | 3.19E-01 |
| *CACNG2* | GDR | 3.12 | 1.54E-02 | 1221 | 0.31 | 1.32E-01 |
| *SNCB* | GDR | 3.11 | 1.56E-02 | 1229 | 0.23 | 3.06E-01 |
| *STXBP5L* | GDR | 3.10 | 1.61E-02 | 1245 | 0.25 | 2.66E-01 |
| *TSPAN9* | GDR | 3.09 | 1.66E-02 | 1259 | 0.26 | 2.34E-01 |
| *ITIH5* | GDR | 3.06 | 1.76E-02 | 1285 | 0.25 | 2.60E-01 |
| *PPP2R2D* | GDR | 3.06 | 1.77E-02 | 1288 | 0.22 | 3.53E-01 |
| *C12orf49* | GDR | 3.05 | 1.83E-02 | 1298 | 0.28 | 1.94E-01 |
| *NDRG4* | GDR | 3.05 | 1.83E-02 | 1300 | 0.23 | 3.04E-01 |
| *SNRNP25* | GDR | 3.04 | 1.84E-02 | 1309 | 0.24 | 2.83E-01 |
| *ECSIT* | GDR | 3.03 | 1.90E-02 | 1330 | 0.28 | 1.89E-01 |
| *SHROOM2* | GDR | 3.01 | 2.01E-02 | 1352 | 0.23 | 3.15E-01 |
| *URM1* | GDR | 2.99 | 2.12E-02 | 1372 | 0.31 | 1.33E-01 |
| *DTX4* | GDR | 2.98 | 2.13E-02 | 1376 | 0.25 | 2.67E-01 |
| *MATK* | GDR | 2.98 | 2.16E-02 | 1383 | 0.28 | 1.94E-01 |
| *TRMU* | GDR | 2.94 | 2.35E-02 | 1427 | 0.29 | 1.77E-01 |
| *CAPN1* | GDR | 2.92 | 2.46E-02 | 1452 | 0.24 | 2.77E-01 |
| *WDR45* | GDR | 2.90 | 2.62E-02 | 1478 | 0.22 | 3.44E-01 |
| *RUSC1* | GDR | 2.87 | 2.77E-02 | 1506 | 0.25 | 2.70E-01 |
| *SEC31B* | GDR | 2.86 | 2.87E-02 | 1524 | 0.22 | 3.36E-01 |
| *MFGE8* | GDR | 2.86 | 2.89E-02 | 1530 | 0.22 | 3.55E-01 |
| *APBA2* | GDR | 2.85 | 2.91E-02 | 1539 | 0.21 | 3.84E-01 |
| *GGA3* | GDR | 2.84 | 3.01E-02 | 1567 | 0.20 | 4.14E-01 |
| *CCDC25* | GDR | 2.83 | 3.07E-02 | 1579 | 0.21 | 3.84E-01 |
| *CHST8* |  | 2.82 | 3.14E-02 | 1591 | 0.29 | 1.65E-01 |
| *MYO1B* | GDR | 2.81 | 3.18E-02 | 1596 | 0.25 | 2.73E-01 |
| *TNFAIP1* | GDR | 2.80 | 3.24E-02 | 1604 | 0.27 | 2.18E-01 |
| *ITM2A* | GDR | 2.80 | 3.25E-02 | 1605 | 0.18 | 4.66E-01 |
| *RPH3A* | GDR | 2.80 | 3.32E-02 | 1611 | 0.26 | 2.45E-01 |
| *CYC1* | GDR | 2.80 | 3.32E-02 | 1612 | 0.23 | 3.04E-01 |
| *SLC24A2* | GDR | 2.79 | 3.33E-02 | 1614 | 0.22 | 3.37E-01 |
| *PENK* | GDR | 2.79 | 3.34E-02 | 1617 | 0.21 | 3.73E-01 |
| *DAPK3* | GDR | 2.79 | 3.38E-02 | 1626 | 0.23 | 3.00E-01 |
| *PMPCA* | GDR | 2.78 | 3.43E-02 | 1631 | 0.26 | 2.34E-01 |
| *FAM49B* | GDR | 2.77 | 3.57E-02 | 1644 | 0.24 | 2.94E-01 |
| *NRSN2* | GDR | 2.76 | 3.62E-02 | 1650 | 0.31 | 1.32E-01 |
| *MTDH* | GDR | 2.75 | 3.68E-02 | 1665 | 0.24 | 2.94E-01 |
| *SPRY2* | GDR | 2.74 | 3.77E-02 | 1684 | 0.20 | 3.95E-01 |
| *FBXL15* | GDR | 2.73 | 3.81E-02 | 1691 | 0.24 | 2.98E-01 |
| *SOX18* | GDR | 2.73 | 3.85E-02 | 1697 | 0.25 | 2.53E-01 |
| *NAPA* | GDR | 2.72 | 3.91E-02 | 1703 | 0.23 | 3.11E-01 |
| *MDH1* | GDR | 2.72 | 3.93E-02 | 1704 | 0.20 | 3.93E-01 |
| *NME1* | GDR | 2.72 | 3.97E-02 | 1712 | 0.19 | 4.43E-01 |
| *ENTPD4* | GDR | 2.71 | 3.99E-02 | 1716 | 0.20 | 4.14E-01 |
| *KATNB1* | GDR | 2.71 | 4.01E-02 | 1723 | 0.21 | 3.67E-01 |
| *GDPD5* | GDR | 2.70 | 4.16E-02 | 1738 | 0.27 | 2.04E-01 |
| *KCNF1* | GDR | 2.69 | 4.21E-02 | 1741 | 0.29 | 1.78E-01 |
| *PPP1R11* | GDR | 2.69 | 4.22E-02 | 1743 | 0.24 | 2.79E-01 |
| *HS3ST2* | GDR | 2.69 | 4.26E-02 | 1747 | 0.28 | 1.85E-01 |
| *EDN3* | GDR | 2.68 | 4.35E-02 | 1763 | 0.18 | 4.54E-01 |
| *POSTN* | GDR | 2.67 | 4.37E-02 | 1768 | 0.18 | 4.56E-01 |
| *KCNJ8* | GDR | 2.67 | 4.37E-02 | 1772 | 0.27 | 2.23E-01 |
| *MPPED1* | GDR | 2.67 | 4.38E-02 | 1776 | 0.25 | 2.57E-01 |
| *C12orf10* | GDR | 2.66 | 4.47E-02 | 1792 | 0.22 | 3.35E-01 |
| *RPP25* | GDR | 2.66 | 4.47E-02 | 1793 | 0.19 | 4.22E-01 |
| *MRPS15* | GDR | 2.66 | 4.52E-02 | 1805 | 0.22 | 3.47E-01 |
| *MGST3* | GDR | 2.65 | 4.58E-02 | 1815 | 0.20 | 4.17E-01 |
| *FSD1* | GDR | 2.65 | 4.61E-02 | 1825 | 0.25 | 2.53E-01 |
| *PLXNA2* | GDR | 2.63 | 4.77E-02 | 1856 | 0.21 | 3.80E-01 |
| *AFTPH* | GDR | 2.62 | 4.83E-02 | 1867 | 0.24 | 2.79E-01 |
| *AIP* | GDR | 2.61 | 4.94E-02 | 1889 | 0.23 | 3.20E-01 |

**Table S5.** List of Gandal Down-Regulated (GDR) genes that were significantly weighted on top PLS1 component. For each gene PLS1, *z*-score represents the bootstrapped *Z*-score of the PLS weight. PLS1 p-value shows the significant level of the PLS1 *z-*score (*z*-test) after FDR-correction. PLS1 ranking shows the position of a gene in the PLS1 *z*-score list. Regional correlation and regional p-value illustrate the pairwise Pearson correlation between gene expression and regional values of schizotypy-related myelination (FDR-corrected).

| **Gene Name** | **Schizophrenia-related list** | **PLS1**  ***Z*-score** | **PLS1**  **Adjusted *P*-value** | **PLS1**  **Ranking out of 20,647** | **Inter-regional**  **correlation** | **Inter-regional**  ***P*-value** |
| --- | --- | --- | --- | --- | --- | --- |
| *SLA* | GUR | -6.11 | 1.16E-06 | 20639 | -0.45 | 1.68E-02 |
| *YPEL1* | GUR | -5.71 | 4.12E-06 | 20622 | -0.41 | 3.41E-02 |
| *EPHX1* | GUR | -5.70 | 4.12E-06 | 20619 | -0.38 | 4.80E-02 |
| *FCGBP* | GUR | -5.57 | 6.90E-06 | 20612 | -0.39 | 4.15E-02 |
| *DPYSL3* | GUR | -5.56 | 6.90E-06 | 20609 | -0.35 | 7.47E-02 |
| *TMEM176B* | GUR | -5.33 | 1.48E-05 | 20581 | -0.37 | 5.68E-02 |
| *TMEM176A* | GUR | -5.30 | 1.61E-05 | 20574 | -0.43 | 2.62E-02 |
| *TLR4* | GUR | -5.22 | 2.17E-05 | 20563 | -0.44 | 2.19E-02 |
| *PEA15* | GUR | -5.19 | 2.40E-05 | 20558 | -0.36 | 7.04E-02 |
| *TTPA* | GUR | -5.15 | 2.54E-05 | 20544 | -0.35 | 7.52E-02 |
| *EDNRB* | GUR | -4.98 | 4.29E-05 | 20497 | -0.32 | 1.08E-01 |
| *PKIA* | GUR | -4.97 | 4.45E-05 | 20495 | -0.38 | 5.08E-02 |
| *ATP2B4* | GUR | -4.97 | 4.57E-05 | 20494 | -0.30 | 1.54E-01 |
| *TRIM24* | GUR | -4.94 | 5.09E-05 | 20490 | -0.48 | 9.96E-03 |
| *POLR2G* | GUR | -4.93 | 5.31E-05 | 20487 | -0.40 | 4.04E-02 |
| *PDLIM5* | GUR | -4.90 | 5.83E-05 | 20480 | -0.35 | 8.35E-02 |
| *PLXDC2* | GUR | -4.89 | 6.15E-05 | 20478 | -0.45 | 1.64E-02 |
| *PEG10* | GUR | -4.83 | 7.53E-05 | 20460 | -0.35 | 7.59E-02 |
| *SFT2D2* | GUR | -4.81 | 8.19E-05 | 20453 | -0.42 | 3.08E-02 |
| *UMPS* | GUR | -4.77 | 9.05E-05 | 20437 | -0.35 | 8.23E-02 |
| *C14orf132* | GUR | -4.76 | 9.18E-05 | 20432 | -0.40 | 3.84E-02 |
| *IL33* | GUR | -4.71 | 1.11E-04 | 20415 | -0.30 | 1.45E-01 |
| *MR1* | GUR | -4.71 | 1.11E-04 | 20414 | -0.38 | 4.80E-02 |
| *DPYD* | GUR | -4.70 | 1.13E-04 | 20411 | -0.36 | 7.33E-02 |
| *SLC1A4* | GUR | -4.64 | 1.44E-04 | 20394 | -0.33 | 1.05E-01 |
| *GPR6* | GUR | -4.62 | 1.53E-04 | 20391 | -0.41 | 3.18E-02 |
| *FCGR1A* | GUR | -4.62 | 1.54E-04 | 20388 | -0.41 | 3.17E-02 |
| *BNIP2* | GUR | -4.57 | 1.87E-04 | 20381 | -0.32 | 1.10E-01 |
| *KCNN3* | GUR | -4.57 | 1.88E-04 | 20377 | -0.29 | 1.73E-01 |
| *FABP7* | GUR | -4.53 | 2.21E-04 | 20367 | -0.34 | 8.75E-02 |
| *GPC4* | GUR | -4.51 | 2.30E-04 | 20361 | -0.34 | 8.87E-02 |
| *APOE* | GUR | -4.49 | 2.46E-04 | 20355 | -0.30 | 1.52E-01 |
| *CD99* | GUR | -4.46 | 2.75E-04 | 20341 | -0.34 | 8.92E-02 |
| *GPM6B* | GUR | -4.42 | 3.18E-04 | 20325 | -0.30 | 1.55E-01 |
| *TRAF3IP2* | GUR | -4.36 | 3.83E-04 | 20302 | -0.29 | 1.69E-01 |
| *TRHR* | GUR | -4.34 | 4.17E-04 | 20292 | -0.31 | 1.27E-01 |
| *CASP6* | GUR | -4.34 | 4.19E-04 | 20291 | -0.35 | 7.84E-02 |
| *ABCA1* | GUR | -4.31 | 4.60E-04 | 20283 | -0.35 | 7.50E-02 |
| *CARHSP1* | GUR | -4.28 | 5.09E-04 | 20273 | -0.31 | 1.38E-01 |
| *CPNE3* | GUR | -4.22 | 6.31E-04 | 20244 | -0.33 | 1.03E-01 |
| *FKBP14* | GUR | -4.20 | 6.67E-04 | 20234 | -0.24 | 2.74E-01 |
| *TAF7L* | GUR | -4.17 | 7.35E-04 | 20217 | -0.45 | 1.85E-02 |
| *FAM65B* | GUR | -4.14 | 8.09E-04 | 20199 | -0.25 | 2.61E-01 |
| *TMEM47* | GUR | -4.13 | 8.15E-04 | 20197 | -0.35 | 8.35E-02 |
| *FAM117A* | GUR | -4.10 | 9.14E-04 | 20182 | -0.28 | 1.82E-01 |
| *DNAJC3* | GUR | -4.10 | 9.14E-04 | 20179 | -0.32 | 1.16E-01 |
| *PLEKHF2* | GUR | -4.07 | 9.85E-04 | 20156 | -0.31 | 1.27E-01 |
| *TRIM27* | GUR | -4.07 | 9.85E-04 | 20155 | -0.29 | 1.60E-01 |
| *HIST2H2BE* | GUR | -4.05 | 1.07E-03 | 20143 | -0.35 | 8.37E-02 |
| *GNG12* | GUR | -4.02 | 1.16E-03 | 20123 | -0.31 | 1.26E-01 |
| *NQO1* | GUR | -3.97 | 1.32E-03 | 20098 | -0.33 | 9.94E-02 |
| *SCN9A* | GUR | -3.97 | 1.32E-03 | 20097 | -0.35 | 8.40E-02 |
| *SSPN* | GUR | -3.93 | 1.53E-03 | 20074 | -0.32 | 1.11E-01 |
| *SLC4A4* | GUR | -3.91 | 1.64E-03 | 20061 | -0.20 | 4.12E-01 |
| *SIDT2* | GUR | -3.90 | 1.65E-03 | 20057 | -0.31 | 1.31E-01 |
| *KAL1* | GUR | -3.90 | 1.65E-03 | 20056 | -0.31 | 1.31E-01 |
| *ERBB2* | GUR | -3.90 | 1.66E-03 | 20054 | -0.32 | 1.17E-01 |
| *GPC5* | GUR | -3.85 | 1.92E-03 | 20013 | -0.28 | 1.91E-01 |
| *SLC7A11* | GUR | -3.84 | 1.97E-03 | 20006 | -0.25 | 2.60E-01 |
| *TMED5* | GUR | -3.80 | 2.21E-03 | 19980 | -0.46 | 1.47E-02 |
| *HSDL2* | GUR | -3.80 | 2.25E-03 | 19976 | -0.29 | 1.72E-01 |
| *CYBRD1* | GUR | -3.77 | 2.44E-03 | 19962 | -0.27 | 2.18E-01 |
| *CETN3* | GUR | -3.76 | 2.49E-03 | 19956 | -0.34 | 8.43E-02 |
| *NPC2* | GUR | -3.75 | 2.59E-03 | 19930 | -0.29 | 1.62E-01 |
| *TTYH1* | GUR | -3.74 | 2.63E-03 | 19927 | -0.28 | 2.00E-01 |
| *AASS* | GUR | -3.73 | 2.69E-03 | 19923 | -0.27 | 2.15E-01 |
| *IFITM2* | GUR | -3.68 | 3.15E-03 | 19869 | -0.38 | 4.80E-02 |
| *TSPAN6* | GUR | -3.66 | 3.28E-03 | 19857 | -0.27 | 2.05E-01 |
| *SMPD2* | GUR | -3.64 | 3.51E-03 | 19835 | -0.24 | 2.94E-01 |
| *ROBO1* | GUR | -3.59 | 4.02E-03 | 19811 | -0.27 | 2.25E-01 |
| *CYP2J2* | GUR | -3.56 | 4.46E-03 | 19791 | -0.22 | 3.37E-01 |
| *ANGEL2* | GUR | -3.53 | 4.91E-03 | 19768 | -0.39 | 4.26E-02 |
| *PPAP2B* | GUR | -3.53 | 4.94E-03 | 19766 | -0.22 | 3.44E-01 |
| *ZZZ3* | GUR | -3.50 | 5.24E-03 | 19746 | -0.33 | 9.91E-02 |
| *ERLIN2* | GUR | -3.47 | 5.70E-03 | 19709 | -0.23 | 3.00E-01 |
| *NECAP2* | GUR | -3.45 | 6.02E-03 | 19675 | -0.23 | 3.00E-01 |
| *MYO10* | GUR | -3.42 | 6.59E-03 | 19651 | -0.20 | 4.04E-01 |
| *HK2* | GUR | -3.41 | 6.75E-03 | 19644 | -0.31 | 1.32E-01 |
| *MPP3* | GUR | -3.40 | 6.79E-03 | 19640 | -0.33 | 9.92E-02 |
| *MCM6* | GUR | -3.40 | 6.84E-03 | 19636 | -0.24 | 2.77E-01 |
| *GSTM2* | GUR | -3.38 | 7.16E-03 | 19618 | -0.21 | 3.72E-01 |
| *ACSS3* | GUR | -3.36 | 7.67E-03 | 19602 | -0.24 | 2.87E-01 |
| *CSTB* | GUR | -3.34 | 8.25E-03 | 19581 | -0.25 | 2.72E-01 |
| *NUPR1* | GUR | -3.33 | 8.44E-03 | 19574 | -0.27 | 2.18E-01 |
| *CFLAR* | GUR | -3.29 | 9.28E-03 | 19541 | -0.30 | 1.58E-01 |
| *GABRB1* | GUR | -3.27 | 9.95E-03 | 19514 | -0.24 | 2.79E-01 |
| *SH3GLB1* | GUR | -3.25 | 1.02E-02 | 19498 | -0.25 | 2.68E-01 |
| *PTPN9* | GUR | -3.25 | 1.03E-02 | 19496 | -0.26 | 2.34E-01 |
| *SELENBP1* | GUR | -3.23 | 1.09E-02 | 19466 | -0.27 | 2.16E-01 |
| *SERP1* | GUR | -3.22 | 1.12E-02 | 19456 | -0.28 | 1.79E-01 |
| *LYN* | GUR | -3.20 | 1.16E-02 | 19438 | -0.27 | 2.09E-01 |
| *HEPH* | GUR | -3.20 | 1.16E-02 | 19436 | -0.17 | 5.18E-01 |
| *TMEM43* | GUR | -3.18 | 1.24E-02 | 19416 | -0.28 | 1.93E-01 |
| *OTUD4* | GUR | -3.18 | 1.24E-02 | 19414 | -0.24 | 2.77E-01 |
| *PTPN13* | GUR | -3.17 | 1.26E-02 | 19405 | -0.25 | 2.61E-01 |
| *HMGA2* | GUR | -3.16 | 1.30E-02 | 19385 | -0.29 | 1.76E-01 |
| *STK17B* | GUR | -3.14 | 1.36E-02 | 19359 | -0.21 | 3.86E-01 |
| *KAT2B* | GUR | -3.13 | 1.39E-02 | 19348 | -0.23 | 3.14E-01 |
| *MT1H* | GUR | -3.13 | 1.40E-02 | 19344 | -0.24 | 2.89E-01 |
| *ZC3HAV1* | GUR | -3.12 | 1.42E-02 | 19335 | -0.26 | 2.47E-01 |
| *PRKG1* | GUR | -3.11 | 1.46E-02 | 19320 | -0.20 | 3.93E-01 |
| *ANGPTL4* | GUR | -3.10 | 1.48E-02 | 19310 | -0.20 | 3.98E-01 |
| *WFS1* | GUR | -3.10 | 1.49E-02 | 19308 | -0.21 | 3.73E-01 |
| *ZNF253* | GUR | -3.10 | 1.49E-02 | 19304 | -0.29 | 1.71E-01 |
| *PLEKHO2* | GUR | -3.07 | 1.59E-02 | 19272 | -0.20 | 4.09E-01 |
| *PDGFRL* | GUR | -3.07 | 1.61E-02 | 19267 | -0.33 | 1.01E-01 |
| *TAF10* | GUR | -3.05 | 1.68E-02 | 19239 | -0.24 | 2.99E-01 |
| *IKZF5* | GUR | -3.05 | 1.69E-02 | 19237 | -0.25 | 2.60E-01 |
| *RARRES3* | GUR | -3.04 | 1.70E-02 | 19232 | -0.23 | 3.08E-01 |
| *BIRC3* | GUR | -3.04 | 1.73E-02 | 19221 | -0.24 | 2.85E-01 |
| *GDPD2* | GUR | -3.03 | 1.74E-02 | 19218 | -0.25 | 2.72E-01 |
| *TBL1X* | GUR | -3.03 | 1.76E-02 | 19214 | -0.26 | 2.45E-01 |
| *ADCYAP1R1* | GUR | -3.03 | 1.77E-02 | 19209 | -0.23 | 3.07E-01 |
| *PAK2* | GUR | -3.01 | 1.83E-02 | 19194 | -0.27 | 2.13E-01 |
| *TIMP1* | GUR | -3.00 | 1.87E-02 | 19180 | -0.23 | 3.00E-01 |
| *CADM1* | GUR | -3.00 | 1.89E-02 | 19176 | -0.22 | 3.55E-01 |
| *ZNF471* | GUR | -2.99 | 1.94E-02 | 19166 | -0.24 | 2.89E-01 |
| *AQP4* | GUR | -2.99 | 1.96E-02 | 19163 | -0.24 | 2.78E-01 |
| *TPP1* | GUR | -2.98 | 2.00E-02 | 19154 | -0.27 | 2.16E-01 |
| *PAICS* | GUR | -2.98 | 2.00E-02 | 19152 | -0.27 | 2.09E-01 |
| *PRSS23* | GUR | -2.98 | 2.01E-02 | 19150 | -0.19 | 4.52E-01 |
| *TMED10* | GUR | -2.98 | 2.01E-02 | 19147 | -0.22 | 3.28E-01 |
| *ZNF350* | GUR | -2.97 | 2.05E-02 | 19128 | -0.20 | 3.96E-01 |
| *SCGB1D2* | GUR | -2.97 | 2.05E-02 | 19127 | -0.22 | 3.45E-01 |
| *CP* | GUR | -2.95 | 2.11E-02 | 19115 | -0.27 | 2.13E-01 |
| *MT2A* | GUR | -2.95 | 2.14E-02 | 19106 | -0.21 | 3.58E-01 |
| *NFE2* | GUR | -2.93 | 2.21E-02 | 19083 | -0.21 | 3.63E-01 |
| *SIRT1* | GUR | -2.93 | 2.23E-02 | 19078 | -0.22 | 3.38E-01 |
| *FAM107A* | GUR | -2.92 | 2.30E-02 | 19060 | -0.19 | 4.22E-01 |
| *SLC1A2* | GUR | -2.91 | 2.31E-02 | 19053 | -0.16 | 5.45E-01 |
| *IGF2BP2* | GUR | -2.91 | 2.34E-02 | 19049 | -0.23 | 3.04E-01 |
| *REST* | GUR | -2.90 | 2.40E-02 | 19042 | -0.23 | 3.23E-01 |
| *CDH20* | GUR | -2.90 | 2.42E-02 | 19038 | -0.25 | 2.60E-01 |
| *STOM* | GUR | -2.89 | 2.43E-02 | 19030 | -0.22 | 3.29E-01 |
| *ITGB8* | GUR | -2.89 | 2.44E-02 | 19029 | -0.16 | 5.56E-01 |
| *C1R* | GUR | -2.88 | 2.54E-02 | 19019 | -0.22 | 3.27E-01 |
| *FOXO1* | GUR | -2.87 | 2.57E-02 | 19013 | -0.27 | 2.06E-01 |
| *YAP1* | GUR | -2.85 | 2.73E-02 | 18987 | -0.20 | 4.05E-01 |
| *ADD3* | GUR | -2.84 | 2.82E-02 | 18973 | -0.27 | 2.22E-01 |
| *GPR56* | GUR | -2.83 | 2.83E-02 | 18968 | -0.19 | 4.43E-01 |
| *DUS4L* | GUR | -2.83 | 2.84E-02 | 18956 | -0.31 | 1.38E-01 |
| *MTHFD2L* | GUR | -2.83 | 2.84E-02 | 18957 | -0.26 | 2.35E-01 |
| *CSGALNACT1* | GUR | -2.82 | 2.89E-02 | 18952 | -0.19 | 4.52E-01 |
| *MLC1* | GUR | -2.82 | 2.89E-02 | 18950 | -0.20 | 4.19E-01 |
| *CTBP2* | GUR | -2.82 | 2.91E-02 | 18947 | -0.17 | 4.92E-01 |
| *ABI1* | GUR | -2.82 | 2.94E-02 | 18939 | -0.20 | 4.00E-01 |
| *CYP7B1* | GUR | -2.81 | 2.95E-02 | 18937 | -0.22 | 3.44E-01 |
| *SERPING1* | GUR | -2.81 | 2.98E-02 | 18935 | -0.31 | 1.39E-01 |
| *ARID1A* | GUR | -2.80 | 3.03E-02 | 18926 | -0.23 | 3.20E-01 |
| *AGT* | GUR | -2.78 | 3.15E-02 | 18895 | -0.28 | 2.00E-01 |
| *METTL7A* | GUR | -2.78 | 3.15E-02 | 18893 | -0.16 | 5.37E-01 |
| *TP53BP2* | GUR | -2.78 | 3.18E-02 | 18887 | -0.19 | 4.38E-01 |
| *FKBP10* | GUR | -2.75 | 3.40E-02 | 18845 | -0.25 | 2.57E-01 |
| *KLF7* | GUR | -2.75 | 3.42E-02 | 18837 | -0.21 | 3.72E-01 |
| *TIMP3* | GUR | -2.75 | 3.43E-02 | 18831 | -0.21 | 3.82E-01 |
| *MT1G* | GUR | -2.74 | 3.48E-02 | 18820 | -0.20 | 4.06E-01 |
| *GSTM4* | GUR | -2.74 | 3.50E-02 | 18817 | -0.23 | 3.14E-01 |
| *NAGA* | GUR | -2.72 | 3.63E-02 | 18786 | -0.23 | 3.12E-01 |
| *NTRK2* | GUR | -2.71 | 3.69E-02 | 18769 | -0.21 | 3.86E-01 |
| *TLR2* | GUR | -2.70 | 3.80E-02 | 18752 | -0.22 | 3.43E-01 |
| *SLC2A5* | GUR | -2.69 | 3.84E-02 | 18737 | -0.23 | 3.02E-01 |
| *PREX2* | GUR | -2.68 | 3.91E-02 | 18720 | -0.24 | 2.91E-01 |
| *RFX3* | GUR | -2.68 | 3.91E-02 | 18714 | -0.21 | 3.56E-01 |
| *SMAD1* | GUR | -2.66 | 4.08E-02 | 18684 | -0.20 | 3.94E-01 |
| *PIR* | GUR | -2.66 | 4.09E-02 | 18681 | -0.20 | 4.06E-01 |
| *TMPRSS5* | GUR | -2.65 | 4.22E-02 | 18665 | -0.19 | 4.23E-01 |
| *ZNF480* | GUR | -2.64 | 4.27E-02 | 18659 | -0.23 | 3.22E-01 |
| *GJA1* | GUR | -2.63 | 4.44E-02 | 18641 | -0.18 | 4.76E-01 |
| *ALDH1L1* | GUR | -2.61 | 4.57E-02 | 18627 | -0.18 | 4.77E-01 |
| *KIAA1598* | GUR | -2.60 | 4.67E-02 | 18613 | -0.22 | 3.37E-01 |
| *BTG1* | GUR | -2.60 | 4.69E-02 | 18611 | -0.28 | 1.87E-01 |
| *CBFB* | GUR | -2.60 | 4.69E-02 | 18608 | -0.21 | 3.63E-01 |
| *ALDH6A1* | GUR | -2.60 | 4.73E-02 | 18603 | -0.20 | 4.00E-01 |
| *H2AFJ* | GUR | -2.59 | 4.77E-02 | 18596 | -0.23 | 3.25E-01 |
| *ROM1* | GUR | -2.59 | 4.77E-02 | 18595 | -0.26 | 2.44E-01 |

**Table S6.** List of Gandal Up-Regulated (GUR) genes that were significantly weighted on bottom PLS1 component. For each gene in PLS1, *z*-score represents the bootstrapped *Z*-score of the PLS weight. PLS1 p-value shows the significance level of the PLS1 *z-*score (*z*-test) after FDR-correction. PLS1 ranking shows the position of the gene in the PLS1 *z*-score list. Regional correlation and regional p-value illustrate the pairwise Pearson correlation between gene expression and regional values of schizotypy-related myelination (FDR-corrected).

### NSPN Consortium member list

**Principal investigators:**

Edward Bullmore (CI from 01/01/2017)

Raymond Dolan

Ian Goodyer (CI until 01/01/2017)

Peter Fonagy

Peter Jones

**NSPN (funded) staff:**

Matilde Vaghi

Michael Moutoussis

Tobias Hauser

Sharon Neufeld

Rafael Romero-Garcia

Michelle St Clair

Kirstie Whitaker

Becky Inkster

Gita Prabhu

Cinly Ooi

Umar Toseeb

Barry Widmer

Junaid Bhatti

Laura Villis

Ayesha Alrumaithi

Sarah Birt

Aislinn Bowler

Kalia Cleridou

Hina Dadabhoy

Emma Davies

Ashlyn Firkins

Sian Granville

Elizabeth Harding

Alexandra Hopkins

Daniel Isaacs

Janchai King

Danae Kokorikou

Christina Maurice

Cleo McIntosh

Jessica Memarzia

Harriet Mills

Ciara O’Donnell

Sara Pantaleone

Jenny Scott

**Affiliated scientists:**

Pasco Fearon

John Suckling

Anne-Laura van Harmelen

Rogier Kievit

Petra Vértes
